# Efficient coding of numbers explains decision bias and noise

**DOI:** 10.1101/2020.02.18.942938

**Authors:** Arthur Prat-Carrabin, Michael Woodford

## Abstract

Human subjects differentially weight different stimuli in averaging tasks, which has been interpreted as reflecting encoding bias. We examine the alternative hypothesis that stimuli are encoded with noise, then optimally decoded. Under a model of efficient coding, the amount of noise should vary across stimuli, and depend on statistics of the stimuli. We investigate these predictions through a task in which participants are asked to compare the averages of two series of numbers, each sampled from a prior distribution that varies across blocks of trials. Subjects encode numbers with a bias and a noise that both depend on the number. Infrequently occurring numbers are encoded with more noise. We show how an efficient-coding, Bayesian-decoding model accounts for these patterns, and best captures subjects’ behaviour. Finally, our results suggest that Wei and Stocker’s “law of human perception”, which relates the bias and variability of sensory estimates, also applies to number cognition.

---

Human choices rely on representations, by the brain, of the variables relevant to decision. How external variables are encoded into internal representations, and how these are then decoded for decision, are central questions in cognitive neuroscience [1–6]. In many decision problems, furthermore, several pieces of information are pertinent; how humans combine these in their decision-making process is a long-standing debate in economics and psychology [7–11]. Recently, a series of experimental studies have focused on averaging tasks, in which subjects are presented with several stimuli (sometimes numbers, but sometimes visual stimuli characterized by their length, orientation, shape, or color) and asked to make a decision about the *average* magnitude of the presented stimuli [12–15]. Although the contribution of each stimulus to the average should, in theory, be proportional to its true magnitude, the weights attributed to stimuli by human subjects, in their decisions, appear to be nonlinear functions of their magnitudes. Subjects asked to compare the averages of two series of digits, for instance, overweight larger digits when making a decision [12]. To account for this seemingly suboptimal behaviour, Refs. [12, 13] show that if the comparison of the average encoded values involves *noise*, then a *nonlinear transformation* of the presented stimuli can partially compensate for the performance loss induced by the noise. The nonlinear distortion of stimuli appears, under this proposal, as a consequence of an optimal encoding strategy, given unavoidable noise at decision time.

Here, we investigate the alternative hypothesis that it is the encoding of a stimulus that is unavoidably noisy, while the decoded value is an optimal estimate of the magnitude that has been encoded by a noisy representation [16, 17]. The resulting noisy estimate will generally be biased, to a degree that will vary as a function of the stimulus [18–21]. Thus the average estimate will be a nonlinear function of the true stimulus magnitude, though for a different reason than in the model of Ref. [12] mentioned above. This nonlinear function will depend on how the encoding noise varies over the stimulus space. Efficient coding theories suggest that the degree of noise with which a stimulus is encoded should be a decreasing function of its probability under the prior distribution of stimuli; in other words, less likely stimuli should be encoded with more noise [5, 22–25]. This implies that the way that both estimation bias and estimation noise vary over the stimulus space should depend on the prior distribution over that space; we test for such dependence in a behavioural experiment.

In a related model that also combines efficient coding with Bayesian decoding, Wei and Stocker derive a relation between estimation noise and estimation bias, a “law of human perception” supported by evidence from numerous sensory domains [26]. In an experiment involving number comparisons, we find support for this law as well, suggesting that it applies to the semantic representation of numerosity carried by Arabic numerals, and not only to the perception of physical stimuli. We also test other implications of both efficient coding and optimal decoding.

These theories highlight the potential role of the prior in human decision-making. Thus we design a task in which subjects are asked to compare the averages of two series of numbers, and manipulate the prior distribution of the presented numbers across different blocks of trials: it is sometimes uniform, but sometimes skewed toward smaller or larger numbers. In each of these three conditions, the weights of numbers in the decisions of subjects appear to vary nonlinearly over the range of presented numbers. We compare the behaviour of our subjects to the predictions of a family of *purely descriptive* statistical models—i.e., models of the distribution of the estimate, conditional on the number presented, with no interpretation in terms of encoding and decoding—, and find that the patterns in their responses are best captured by a model in which a nonlinear transformation of a presented number is observed with a degree of noise that itself depends on the number.

We then turn to a two-stage *encoding-decoding* model—i.e., a model that decomposes the estimation process into an encoding stage and a decoding stage, each of which is further assumed to be optimized for the distribution of stimuli encountered in the task. We show how a model of efficient coding combined with Bayesian decoding accounts parsimoniously both for the nonlinear transformation and for the noise, and best captures subjects’ behaviour. We find that the encoding noise varies depending on the prior, and that this allows our subjects to increase their expected reward in the task.

## Results

We first present our average-comparison experiment. In each trial of a computer-based task, ten numbers, each within the range [10.00, 99.99], and alternating in color between red and green, are presented to the subject in rapid succession. The subject is then asked to choose whether the five red numbers or the five green numbers had the higher average (Fig. 1a). In different blocks of trials, numbers are sampled from different prior distributions: the **“Uniform”** distribution, under which all numbers in the [10.00, 99.99] range are equally likely, an **“Upward”** distribution (under which higher numbers are more likely), and a **“Downward”** distribution (under which lower numbers are more likely) (Fig. 1b). Within a block of trials, both red and green numbers are sampled from the same distribution. Additional details on the task are provided in Methods. In our analyses, below, we combine the responses of all the subjects; the examination of individual data further supports our results (see Methods).

**Figure 1:**
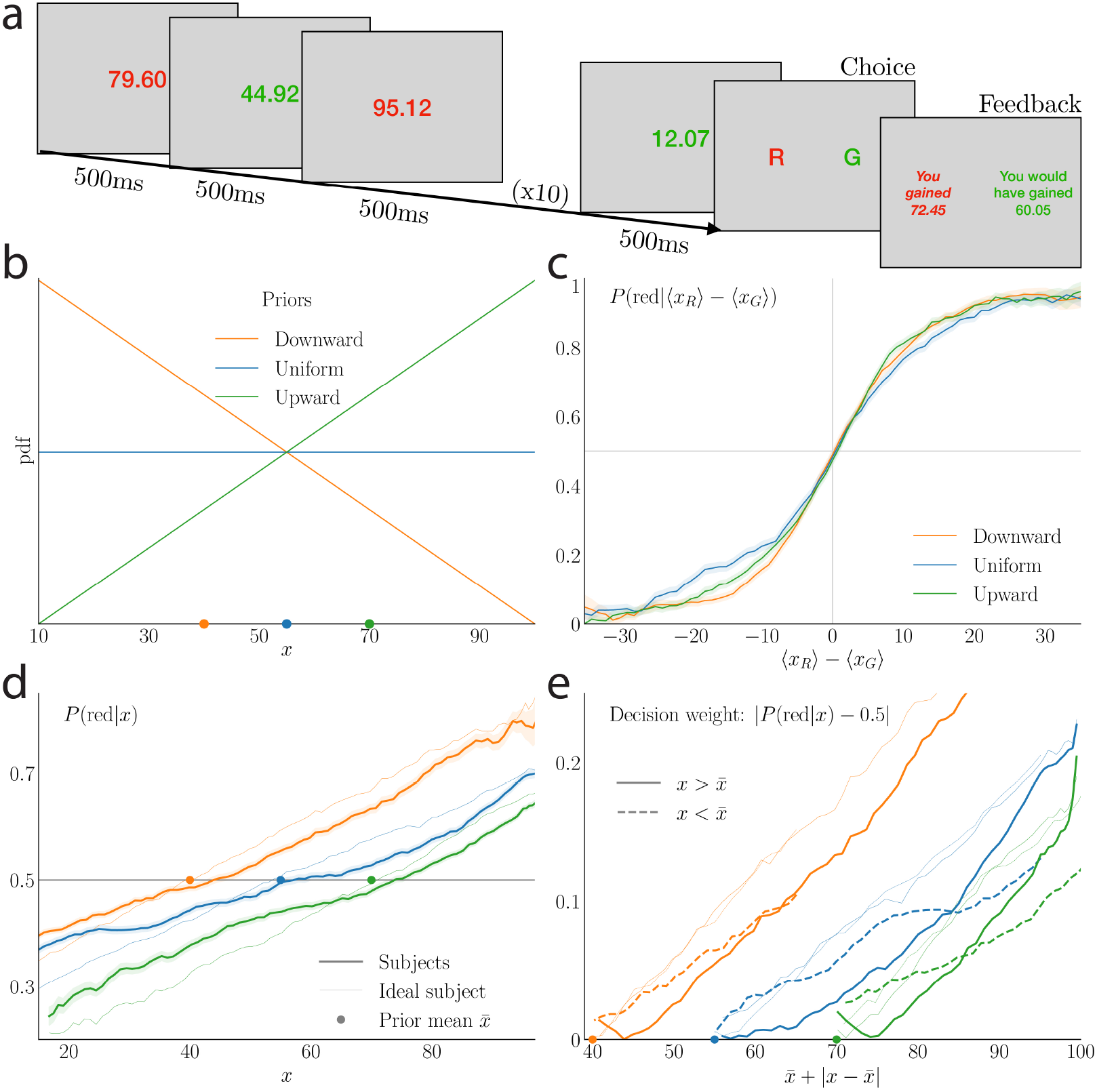
Illustration of the task and subjects choice behaviour. **a**. In each trial, ten numbers are sequentially presented, alternating in color between red and green. The subject can then choose a color, without time constraint, by pressing the corresponding key. Feedback is given, which reports the two averages and emphasizes the chosen one. **b**. In each trial, both red and green numbers are drawn from the same prior distribution over the range [10.00, 99.99]. In different blocks of consecutive trials, the prior is either Downward (triangular distribution with the peak at 10.00, orange line), Uniform (blue line), or Upward (triangular distribution with the peak at 99.99, green line). **c**. Choice probability: in each of the three prior conditions, the fraction of trials in which subjects choose the ‘red’ average, as a function of the true difference in the averages of the two series of numbers. While an ideal subject would be described by a step function, the choices of subjects result in sigmoid curves. **d**. Probability *P*(red|*x*) of choosing ‘red’ conditional on a red number *x* being presented, as a function of *x*, for the subjects (thick lines) and the ideal subject (thin lines). **e**. Decision weight, defined as |*P*(red|*x*) −0.5 |, for the subjects (thick lines) and the ideal subject (thin lines), as a function of the sum of the prior mean, 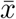, and the absolute difference between the number and the prior mean, 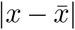. Subjects’ decision weights for numbers below the mean (dashed line) and above the mean (solid line) are appreciably different, whereas for the ideal subject there is only a small difference due to sampling error. In **c, d**, and **e**, each point of the curves is obtained by taking the average, combining all subjects’ data, of the quantity of interest (ordinate) over a sliding window of length 10 of the quantity on the abscissa, incremented by steps of length 1. In **c** and **d**, the shaded areas represent the standard error of the mean.

Although the presented information would allow, in principle, an unambiguous identification of the color of the numbers that have the higher average, subjects make errors in their responses. These errors are not independent of the presented numbers: the proportion of trials in which they choose ‘red’ (the “choice probability”) is an increasing, sigmoid-like function of the difference between the average of the red numbers and that of the green numbers, with about 50% red responses when this difference is close to zero. In other words, subjects make less errors when there is a marked difference between the two averages (Fig. 1c).

In order to isolate the influence of a single number on the decision, we can compute the probability of choosing ‘red’ conditional on a given (red) number *x* being presented, *P* (‘red’|*x*), and the absolute difference between this probability and 0.5: |*P* (‘red’ | *x*) −0.5 |. This quantity, which we call the “decision weight” (following Ref. [12]), is a measure of the average impact of a given number on the decision. For an ideal subject who makes no errors, the decision weight should be zero for *x* equal to the prior mean 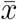, and should increase approximately linearly with the absolute distance 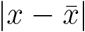 from the prior mean (Fig. 1d,e, thin lines). In our subjects’ data, instead, the value of *x* for which the decision weight is zero is somewhat larger than 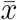 for each prior, and the decision weight increases more steeply with increases in *x* above this value than it does with decreases in *x* below that value (Fig. 1d,e, thick lines).

We wish to explore models of noisy comparison that can account for these regularities. We begin by fitting models to our data that do not separate the processes of encoding and decoding, and instead simply posit that a given number *x* results in a noisy estimate 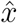 drawn from a conditional distribution 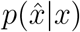. We assume that the ten numbers presented are estimated with this procedure (with independent draws), and that the color ‘red’ is chosen if the average of the estimates of the red numbers is greater than that of the green numbers, i.e., if 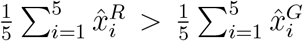, where 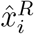 and 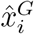 are the estimates for the red and green numbers. Note that this kind of characterization of the data is equally consistent with the theory of Ref. [12], in which 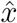 is equal to a deterministic transformation of *x* plus a random noise term added at the time of the comparison process, and a model in which *x* is encoded with noise and 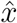 is then an estimate of the value of *x* based on the noisy encoded value.

We consider several possible assumptions about the form of the conditional distribution of estimates,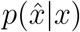. The simplest kind of model consistent with a sigmoid curve of the kind shown in Figure 1c would be one in which the estimate is normally distributed around the true value of the number, with a constant standard deviation, *s*, i.e.: 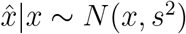. In other words, numbers are perceived with constant Gaussian noise, but the estimates are unbiased. The probability of choosing the red color, conditioned on the ten presented numbers, would then be

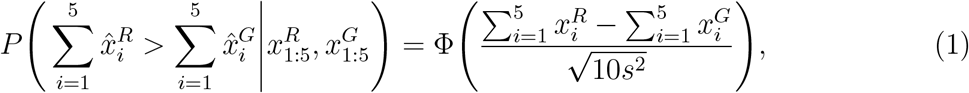

where Φ is the cumulative distribution function (CDF) of the standard normal distribution. The choice probability in this model is thus a sigmoid function of the difference between the averages of red and green numbers, similarly to the choice probability of subjects (Fig. 2a). However, the decision weight of a number, in this model, is an approximately linear function of the absolute difference between the number and the prior mean, and two numbers equally distant from the mean have the same weight (Fig. 2c, top left panel). Thus this simple model does not reproduce the subjects’ unequal weighting of numbers in decisions documented in Figure 1e.

**Figure 2:**
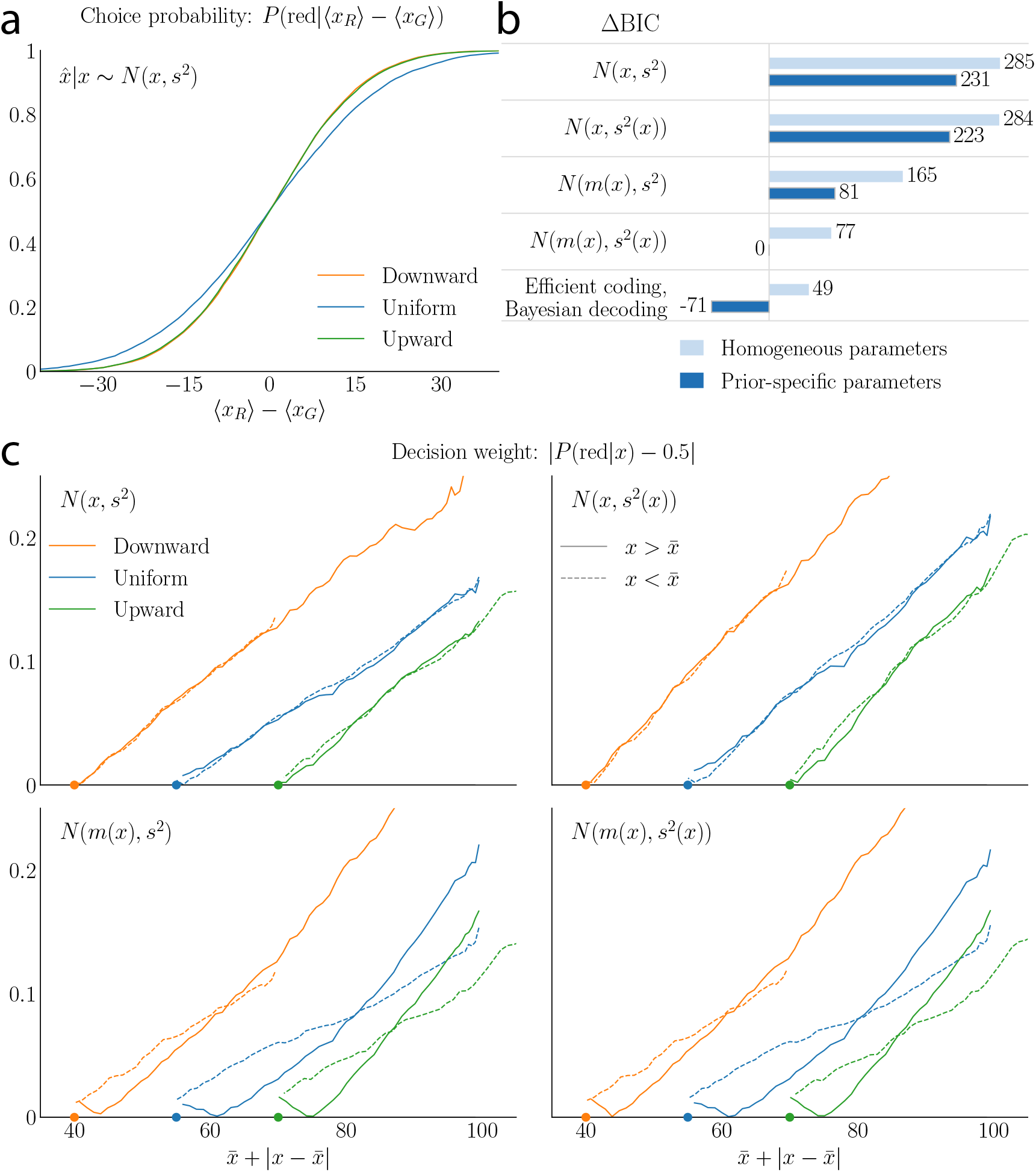
Models with both nonlinear transformation and varying noise best capture subjects’ behaviour. **a**. Choice probability under the unbiased, constant-noise model (*N* (*x, s*^2^)) as a function of the difference in the averages of the presented numbers, for the three prior conditions. **b**. Model comparison statistics for the four one-stage models of noisy estimation and for the efficient-coding, Bayesian-decoding model, both with homogeneous parameters and with prior-specific parameters. For each model class, the difference in BIC from the least restrictive model class (general transformation, variable noise, prior-specific) is reported. The lower BIC for the efficient-coding, Bayesian-decoding model (a special case of the general one-stage model) indicates that it captures subjects’ behaviour more parsimoniously. **c**. Decision weights |*P*(red|*x*) − 0.5| for the unbiased, constant-noise model (top left), the unbiased, varying-noise model (top right), the model with transformation of the number and constant noise (bottom left), and the model with transformation of the number and varying noise (bottom right). Only the two models with transformation of the number reproduce the behaviour of subjects shown in Fig. 1e. All models are fit to the subjects’ combined data.

In the model just presented, estimates are unbiased 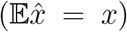, and all numbers are estimated with the same amount of noise 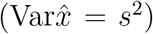. But we also consider models that are more flexible in either or both of these respects. For example, we can allow for bias by assuming that the estimate, 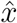, of a presented number, *x*, is normally distributed around a nonlinear transformation, *m*(*x*), of the number: 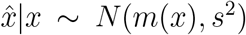. Such a model is equivalent to the one assumed in Refs. [12, 13], in which noise is added to the sum of nonlinearly transformed values of the stimuli. Alternatively, we might assume that the variability of estimates is not constant over the stimulus space, as is found to be true in many stimulus domains [20]. The simplest case of this kind would be one in which 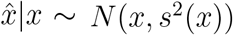, where now the estimation noise, *s*(*x*), varies with the number. Our most general model combines the features of these last two, positing that a transformation of the number is observed with varying noise: 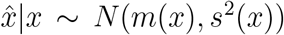. In this model, the probability of choosing the red color, conditioned on the ten presented numbers, will be

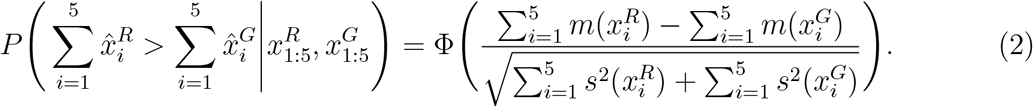

The first three models are special cases of the fourth one, with either an absence of bias (i.e., an identity transformation *m*(*x*) = *x*), a constant noise (*s*(*x*) = *s*), or both (Eq. (1)).

We fit these models to the behavioural data by maximizing their likelihoods. We flexibly specify the functions *m*(*x*) and *s*(*x*), by allowing them to be arbitrary low-order polynomials, defined by their values at a small number of points (between 2 and 8), with the order chosen to minimize the Bayesian Information Criterion (BIC). Running simulations of the fitted models, using the numbers presented to the subjects, we find that each of them predicts that choice frequency should be a similar sigmoid function of the difference between the two averages. The models differ, however, in the implied graphs for the decision weights. The unbiased model with varying noise (*N* (*x, s*^2^(*x*))) still results in approximately linear decision weights, with equal weights for numbers at equal distances from the mean (Fig. 2c, top right panel). Instead the two models that include a transformation *m*(*x*) of the presented numbers yield nonlinear decision weights resembling those of the subjects: numbers below the mean and close to it have higher weights that numbers above the mean and equally distant from it, while the opposite occurs for numbers further from the mean (Fig. 2c, bottom panels. More details on how the transformation *m*(*x*) affects the decision weights can be found in Methods.) In summary, the two models which feature a transformation of the number, whether with a constant or variable noise, better capture the patterns observed in behavioural data.

In order to quantitatively compare the degree of explanatory success of the models, we compute the BIC for the best-fitting model in each class. We can either assume that the parameters of the models are identical under the three priors (“homogeneous parameters”), or, conversely, that the parameters depend on the prior (“prior-specific parameters”). The latter results in a larger number of parameters, which is penalized by the BIC. In spite of this penalty, in all four models the BIC is lower with prior-specific parameters than with homogeneous parameters, suggesting that subjects’ decision-making process is adapted to the prior distribution (Fig. 2b). The two best-fitting one-stage models include a transformation *m*(*x*) of the presented number, in accordance with our observations about the decision weights, and the best-fitting model also features a variable noise *s*(*x*). We note that the BIC in this case is significantly lower than with a transformation but maintaining constant noise (ΔBIC = 81), in spite of the additional parameters of the noise function *s*(*x*). In Methods we discuss the finer features of subjects’ behaviour that are better captured by allowing for variable noise, showing that a nonlinear transformation alone (as in Ref. [12, 13]) does not suffice to account for our data. In addition, to investigate the robustness of our results, we conduct a random-effect analysis that takes into account the subjects’ heterogeneity. We follow the Bayesian model selection procedure described in Ref. [27], and use cross-validation to estimate the models’ likelihoods. This analysis, detailed in Methods, supports the results presented here.

The shape of the best-fitting noise function *s*(*x*) for each prior is shown in Figure 3a. In each case, we find that it is U-shaped: minimal at a value close to the mean of the prior (but slightly below it), and increasing as a function of the distance from this minimum. In the Downward-prior condition, larger numbers (which have a low probability) are perceived with a much larger amount of noise than small numbers (which have a high probability). The noise function in the Upward condition approximately mirrors that in the Downward condition, with small numbers perceived with more noise than large numbers. In other words, the least likely numbers are perceived with the greatest randomness, suggesting that the process by which subjects estimate numbers is adapted to the prior distribution from which they are drawn.

**Figure 3:**
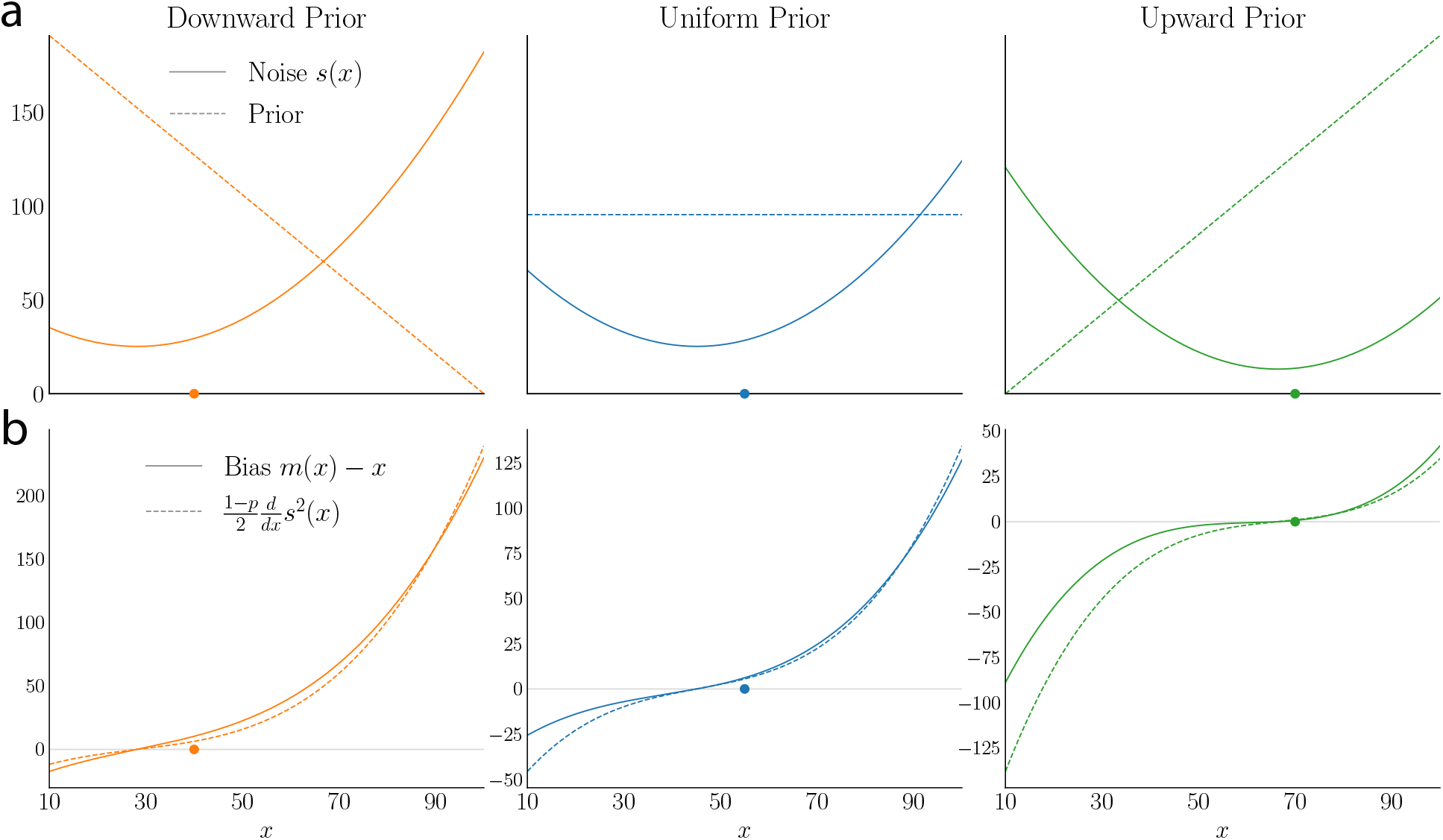
Best-fitting noise and bias: subjects encode less frequent numbers with greater noise. **a**. The best-fitting noise function *s*(*x*) in the *N* (*m*(*x*), *s*^2^(*x*)) model (solid line), and prior distribution (dashed line), in the Downward (left), Uniform (middle) and Upward (right) conditions. The scale refers to the noise (scale of the prior pdf not shown.) **b**. The bias (solid line) and the derivative of the noise variance (multiplied by 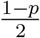, dashed line) for the best-fitting *N*(*m*(*x*), *s*^2^(*x*)) model, as a function of *x*, in the three prior conditions. The efficient-coding, Bayesian-decoding model requires these two functions to be the same (Eq. (4)). Model fit to the subjects’ combined data.

The finding that the variance of 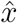 differs for different numbers *x* is a natural one if we suppose that each estimate 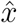 results from an encoding-decoding process, whereby each individual number is encoded, with noise, into an internal representation *r*, and the estimate 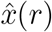 represents an *inference* about the likely value of *x*, based on its noisy internal representation *r*. In general, an optimal rule of inference will make the estimate 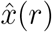 a nonlinear function of *r*, so that the variance of 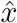 should depend on *x* even if we suppose that the variance of *r* does not. At the same time, an encoding-decoding model of this kind, in which decoding is assumed to be optimal given the nature of the noisy encoding, will imply that the function *m*(*x*), indicating the bias in the average decoded value, will not be independent of *s*(*x*). Thus this class of models represents a special case of the general specification *N* (*m*(*x*), *s*^2^(*x*)), though a different restricted class than any of those considered above. In short, while hitherto we have investigated several assumptions about the form of the conditional distribution of estimates, 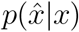, we now derive this distribution from a more parsimonious model of encoding-decoding, and examine its ability to capture the behavioral data.

The model that we consider, based on a proposal of Morais and Pillow [28] that we further discuss in Ref. [29], combines a noisy but efficient coding of the presented number, with a Bayesian-mean decoding of the noisy internal representation. More precisely, suppose that a presented number, *x*, elicits in the brain a series of *n* signals, *r* = (*r*_1_, …, *r*_*n*_), each drawn independently from a distribution conditioned on the presented number, *p*(*r*_*i*_ |*x*). The Fisher information, *I*(*x*), of the vector *r* is *n* times that of one individual signal, *r*_*i*_. Suppose that this encoding is efficient, in the sense that *I*(*x*) solves the optimum problem

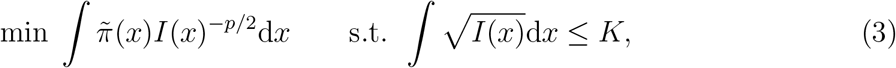

where *p* > 0, and where 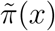 is a subjective prior about the distribution of numbers, which may differ from the prior actually used in the experiment, *π*(*x*). The constraint in Eq. (3) is the same as in Ref. [30], though we consider a more general objective. The rationale for minimizing the objective assumed in Eq. (3) is that, as shown in Ref. [28], it provides a lower bound on the mean *L*_*p*_ error of any point estimator of *x*. In the limiting case *p* → 0, minimizing this quantity is equivalent to maximizing an approximation of the mutual information between the number and its internal representation, which is the objective assumed by Wei and Stocker [30]. An objective with *p* > 0 is more plausible, however, in our setting, because in our task, larger estimation errors reduce a subject’s reward to a greater extent than small errors do. The optimal Fisher information, derived from Eq. (3) through variational calculus [28, 29], is proportional to a power of the subjective prior: 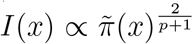 (see Methods).

As for the decoding, we assume that the estimate of subjects, 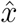, is the mean of the Bayesian posterior over the numbers, *x*, given the noisy signal, *r*. With large *n*, this decoding rule, combined with the efficient coding scheme just presented, implies asymptotically a relation between the bias of the estimate and the derivative of the inverse Fisher information [29] (see Methods), as

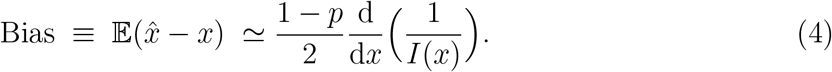

The estimate variance, in addition, is approximately 1*/I*(*x*). Thus we approximate the distribution of estimates as

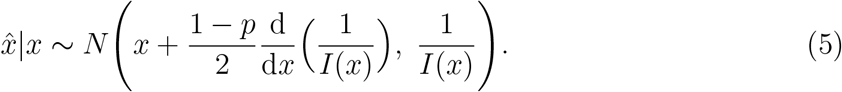

(These asymptotic approximations to the bias and to the variance are accurate to the order 1*/n*; see Methods.) As in the one-stage models considered above, the estimate is normally distributed around a transformation of the number. This is a special case of the general model considered above (*N*(*m*(*x*), *s*^2^(*x*))), with the added constraint that both the nonlinear transformation *m*(*x*) and the noise *s*(*x*) are here determined by a single function, the Fisher information *I*(*x*). In particular, the efficient-coding, Bayesian-decoding model predicts that the bias, *m*(*x*) − *x*, is proportional to the derivative of the noise variance, *s*^2^(*x*) = 1*/I*(*x*) (Eq. (4)).

We can test whether our data are consistent with the additional restriction given by Eq. (4). In Figure 3b, we plot the functions corresponding to the two sides of this equation, implied by our best-fitting estimates of *m*(*x*) and *s*(*x*) for each of the three conditions, when the theoretical restrictions implied by the efficient-coding, Bayesian-decoding model are *not* imposed in the estimation. Here it is important to note that the transformation *m*(*x*) can be identified from choice data only up to an arbitrary affine transformation; this gives us two free parameters to choose in plotting the left-hand side of the equation. As explained in Methods, we choose these parameters to make the implied function more similar to the right-hand side of the equation. When we do so, we obtain the functions plotted in Figure 3b. As noted above, the noise *s*(*x*) is U-shaped; thus the derivative of the noise variance, 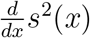, increases as a function of *x*; it vanishes at a value close to the prior mean, is negative below this value, and is positive above it (Fig. 3b, dashed line). With an appropriate choice of normalization for *m*(*x*), the bias vanishes at the same value; in addition, we note that for all three priors the bias is also negative below this value and positive above it, and increases as a function of *x* (Fig. 3b, solid line). In other words, numbers below the prior mean are underestimated, while numbers above the prior mean are overestimated. Thus we find that the two functions, when fitted to subjects’ data, are qualitatively consistent with the predicted relation (Eq. (4)).

We can also estimate directly the model of estimation bias implied by Eq. (5), finding the parameter *p* and the Fisher information function *I*(*x*) that minimize the BIC. (Here also, we allow the noise function to be a low-order polynomial; in Methods, we consider a different functional form, closer to that used in Ref. [12], with similar results.) We assume that, regardless of the distribution of numbers, the subjects optimize the same efficient coding criterion, determined by *p*; thus even when fitting with prior-specific parameters, we maintain the parameter *p* constant across conditions. As with our one-stage models, prior-specific parameters yield a lower BIC than homogeneous ones. Furthermore, the BIC is lower than that of the model in which the functions *m*(*x*) and *s*(*x*) are unrestricted (Fig. 2b, ΔBIC = 71). We conclude that the model implied by Eq. (5) captures more parsimoniously the behaviour of subjects.

In this model, a single function, the Fisher information, *I*(*x*), or equivalently the noise function, 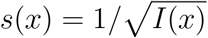, together with the value of *p* completely determines the statistics of responses. Moreover, the assumption that the encoding is optimized, in our model (Eq. (3)), results in the prediction that the noise function should be proportional to the inverse of the subjective prior, 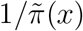, raised to the power 1*/*(*p* + 1). As we do not have access to the subjective prior, we show, in Figure 4, the noise functions alongside the correct priors raised to the power −1*/*(*p* + 1), with *p* equal to its best-fitting value, 0.7. The best-fitting noise function differs under the three priors. In the Downward condition, the noise is an increasing function of the number over most of the range of presented numbers (Fig. 4, left panel). In the Upward condition, conversely, the fitted noise function decreases with the number (except at large numbers; Fig. 4, right panel). Hence, in both the Downward and Upward conditions, the noise function is higher for numbers that are less likely under the prior. In the Uniform condition, the noise reaches a minimum around 40, and large numbers are observed with somewhat more noise than small numbers (Fig. 4, middle panel).

**Figure 4:**
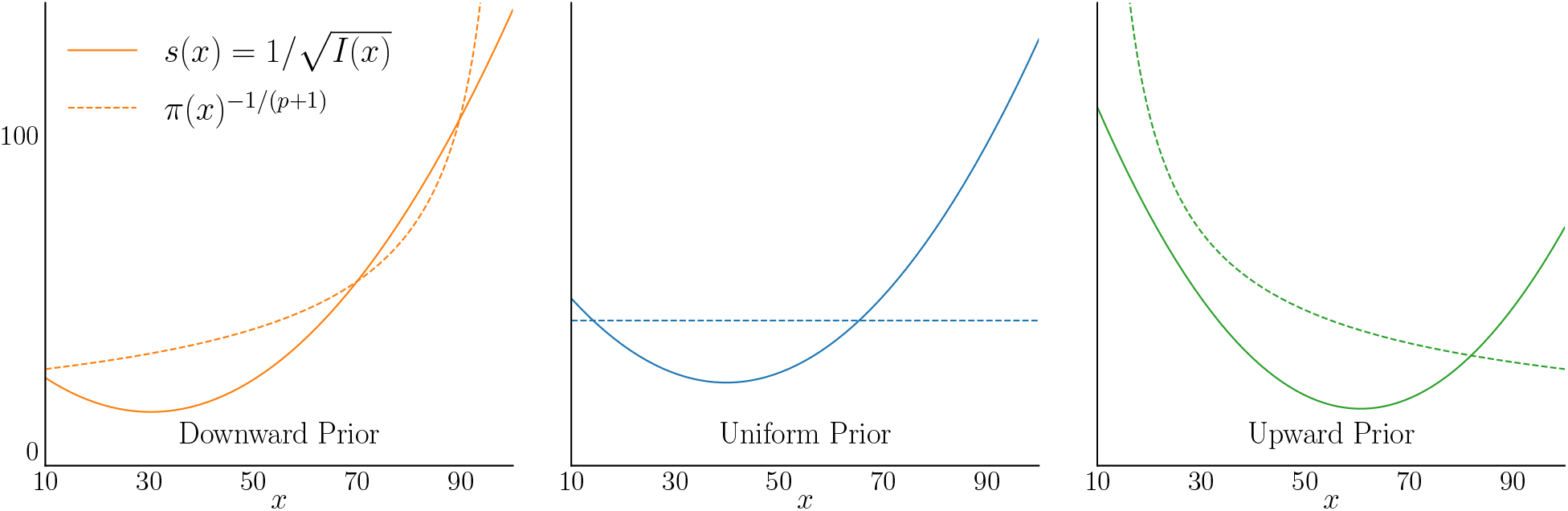
Efficient-coding, Bayesian-decoding model: the Fisher information, fitted to subjects’ data, is adapted to the prior. Noise function, *s*(*x*), equal to the inverse square-root of the Fisher information, fitted to subjects’ data (solid lines), and prior *π*(*x*) raised to the power −1*/*(*p* + 1), with the best-fitting value *p* = 0.7 (dashed lines), in the Downward (left), Uniform (middle) and Upward (right) conditions. The ordinate scale refers to the noise function (scale for the prior not shown). Model fit to the subjects’ combined data.

The behavioral implications of these fitted functions are shown in Figure 5. The estimation bias is proportional to the derivative of the squared noise function (Eq. (4)): it vanishes where the noise reaches a minimum; for smaller *x*, it is negative, while for higher *x* it is positive. Moreover, the efficient-coding, Bayesian decoding model reproduces the unequal weighting of numbers in subjects’ decisions (Fig. 5c,d) and the sigmoid shape of the choice probability curve (Fig. 5e).

**Figure 5:**
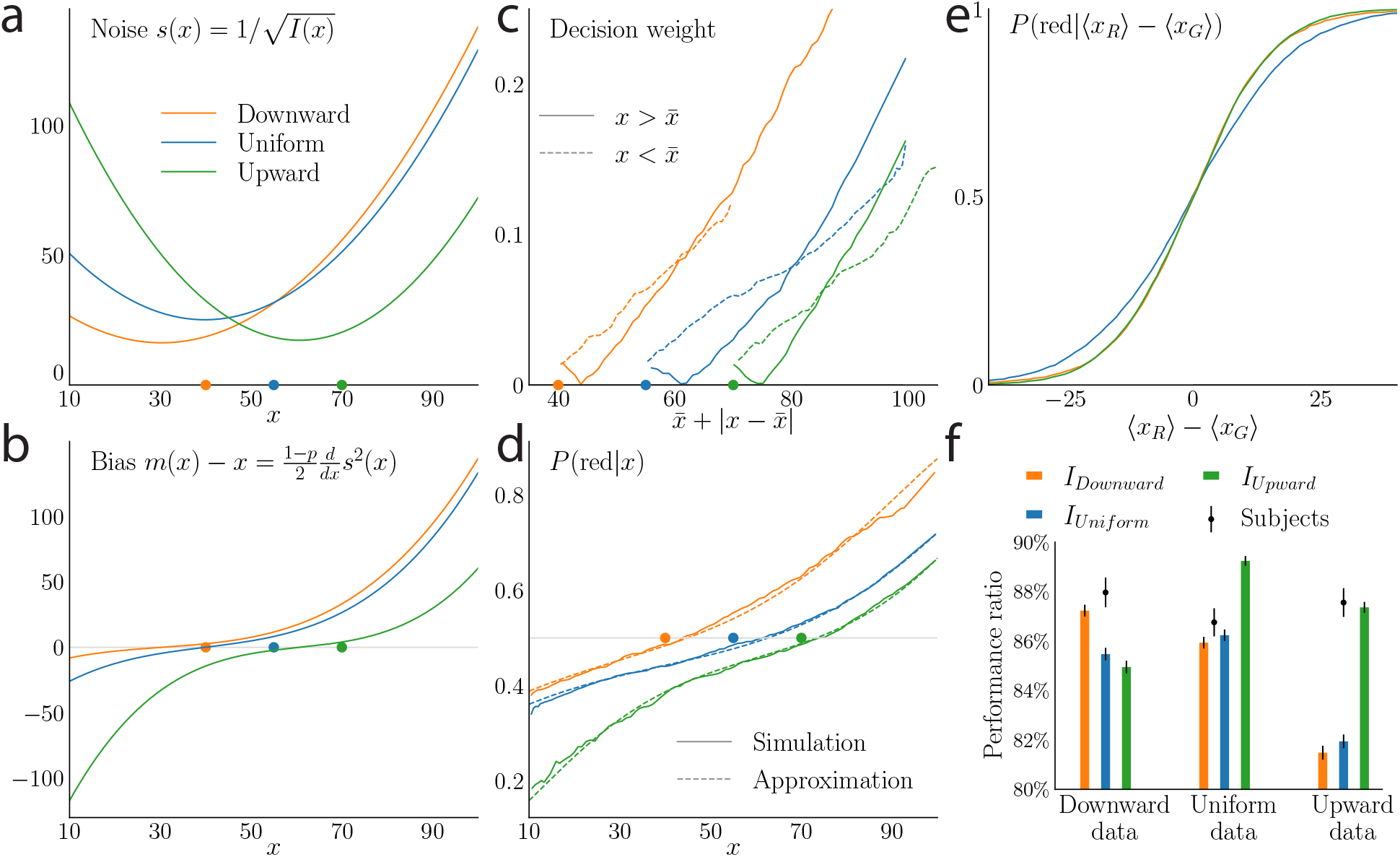
The efficient-coding, Bayesian decoding model reproduces subjects’ behaviour, and the Fisher information functions fitted to subjects’ data improve the performance ratio, in the Upward and Downward conditions. **a**. Noise *s*(*x*) implied by the Fisher information, in the three prior conditions. **b**. Bias *m*(*x*) − *x* in the three prior conditions, also implied by the Fisher information. **c**. Decision weights |*P*(red|*x*) − 0.5 |predicted by the efficient-coding, Bayesian decoding model. **d**. Probability of choosing ‘red’ conditional on a red number *x* being presented, in the model (solid line), and our approximation to the model prediction (dashed line; see Methods). **e**. Choice probabilities implied by the model as a function of the difference between the averages of the numbers presented. **f**. Performance ratios, over numbers sampled from the Downward (left), Uniform (middle) and Upward (right) priors, for the subjects (black dots) and implied by the estimated encoding rules (bars) for each of the three prior conditions. With numbers sampled from the Downward prior, the Downward encoding rule, *I*_*Downward*_, yields the best performance (left orange bar), whereas with numbers sampled from the Upward prior, the Upward encoding rule, *I*_*Upward*_, results in the best performance (right green bar). The black vertical lines represent the standard deviations of the performance ratios (for the subjects, this quantity is bootstrap-estimated from the data). All analyses use the subjects’ combined data.

Our fitted model implies substantial estimation biases, in particular for the less probable numbers (Fig. 5b). In our model, these biases can exist only to the extent that noise is both substantial in magnitude and sufficiently variable over the range of numbers used; however, the degree of noise required is consistent with the variability of our subjects’ responses. Moreover, it is not implausible that our subjects should encode individual numbers with considerable imprecision, given that they must process ten numbers, each presented for only 500 ms.

The theoretical predictions shown in Figure 5 are based on a small-noise approximation, which need not be accurate when noise is substantial. Numerical simulations indicate that the bias implied by the small-noise approximation is greatest in the case of extreme values of *x* that occur with low prior probability, and thus it seems likely that the size of the predicted bias for extreme values of *x* is exaggerated in Fig. 5b. However, because these values do not often occur in our sample, we do not believe that our conclusions regarding the degree of fit of the Bayesian decoding model are greatly influenced by this approximation error; see Methods for a detailed analysis.

Our estimates best fit subjects’ data (even penalizing the additional free parameters) when the Fisher information is allowed to differ depending on the prior distribution from which numbers are sampled. Context-dependent encoding of this kind is predicted by theories of efficient coding; this leads us to ask whether the differing encoding rules that we observe represent efficient adaptations to the different priors, in the sense of maximizing average reward in our task. We investigate this hypothesis by examining the performance, over numbers sampled from a given prior (say, Downward), of a model subject equipped with the Fisher information fitted to subjects’ data in the context of another prior (say, Upward). In other words, we look at how successful the “Upward encoding” (and associated Bayesian decoding) would be on “Downward data” (we use these shorthands below). We measure performance as follows. In each trial of our task, the score is augmented by the chosen average, so that the subject is guaranteed to receive at least the minimum of the two averages. The additional value to be captured in a trial *i* is thus the absolute difference between the two, i.e., Δ_*i*_ = |⟨*x*^*R*^⟩_*i*_ − ⟨*x*^*G*^⟩_*i*_|. We define the “Performance ratio” as the fraction of this value captured by a model subject, i.e., ∑*δ*_*i*_Δ_*i*_/∑Δ_*i*_, where *δ*_*i*_ is 1 if the model subject makes the correct choice in trial *i* and 0 otherwise.

With Downward data, the Downward encoding yields the largest performance ratio (87.2%, s.d.: 0.47%), whereas the Upward encoding results in the lowest ratio (84.9%, s.d.: 0.51%, Fig. 5f, left bars). Conversely, with Upward data the Upward encoding outperforms the Downward encoding (87.4% vs. 81.5%, s.d.: 0.44% and 0.56%, Fig. 5f, right bars). In other words, the encoding rule fitted to the behaviour of subjects in the context of a given prior, Upward or Downward, results in higher rewards in the context of numbers sampled from this same prior, suggesting that the choice of encoding rule is efficient. With Uniform data, the performance ratio of the Uniform encoding is slightly better than that of the Downward encoding rule, but not quite as good as that of the Upward encoding rule (86.2% vs. 85.9% and 89.2%, sd: 0.47%, 0.47%, and 0.39%). However, the performance ratio of the subjects in the Uniform condition (86.8%, s.d.: 1.1%) is lower than that in the Upward (87.6%, s.d.: 1.2%) condition. Consistent with this, the fitted Uniform encoding rule is noisier than the Upward one; and it is primarily this fact (rather than the way that the precision of encoding varies for different numbers) that makes the fitted Uniform encoding less efficient even for Uniform data.

## Discussion

We designed an average-comparison task in which we changed the prior distribution from which numbers were sampled across blocks of trials. In their choices, subjects seem to differentially weight numbers that should be equally relevant to the correct decision. Subjects’ behaviour can be characterized by a model of noisy perception of the size of numbers, in which both the average estimation bias and the variability of estimates depends on the magnitude of the number. Furthermore, we introduced an encoding-decoding model, in which the variable precision of encoding is specified by a Fisher information function that is efficiently chosen with regard to a subjective prior, and the decoded value of each number is the Bayesian-mean estimate based on the encoded evidence. This is equivalent to a constrained version of the model of noisy perception (Eq. (5)), in which both the bias and the variable noise are determined by a single function, the Fisher information. This implies a relation between the bias and the derivative of the noise variance (Eq. (4)), for which we find qualitative support in data. The efficient-coding, Bayesian-decoding model yields the lowest BIC among the models considered, and reproduces the patterns observed in behaviour. Furthermore, the Fisher information fitted to subjects’ data varies depending on the prior distribution. With the two skewed priors (Upward and Downward priors,) the Fisher information is lower for numbers less likely to appear under the prior (Fig. 4). This resulted in a higher performance in the task (Fig. 5f), suggesting that subjects efficiently adapted their decision-making process to the prior distribution of presented numbers. Our results are supported by a Bayesian random-effect analysis, which takes into account the subjects’ heterogeneity (see Methods).

Reference [12] had previously found that a model featuring a nonlinear transformation of presented numbers (i.e., a bias) reproduced the apparent unequal weighting of numbers in subjects’ decisions, and interpreted the bias as a strategy to compensate for decision noise. We provide a different account, that relies on two further observations. First, the noise in the perceived value of a number seems to vary with the number, and to depend on the prior (Fig. 3a; see also the Methods, in which we exhibit behavioral patterns that are captured by the models featuring both nonlinear transformation and varying noise, but not by the model that features a nonlinear transformation only). Second, the way that the bias and the noise function vary from one condition to another is consistent with the theoretical relation between these two functions predicted by our model of efficient coding and Bayesian decoding. Under this account, estimation noise, estimation bias, and thus the unequal weighting of numbers in average comparisons and the sigmoid-shape choice probability function, all derive from the form of the Fisher information function. In turn, the Fisher information is an increasing function, at least roughly, of the prior density from which numbers are sampled, suggesting that the encoding is efficiently adapted to the prior.

We do not, however, reject the hypothesis that there may also be noise in the decision process. Our models, in fact, are not inconsistent with this hypothesis. Decision noise could be included in our models by adding a noise term to the decision variable (defined as the difference between the estimated empirical averages), as in Ref. [12]. This would add a constant to the expression inside the square-root sign in Eq. (2), but the same effect would result from adding a constant to the function *s*^2^(*x*) in our model. The noise functions that we fit can be understood as implicitly already including this constant; note that if we assume that *s*^2^(*x*) is equal to 1*/I*(*x*) plus a constant (rather than simply 1*/I*(*x*), as stated above), the relationship between the bias and the variance of estimates implied by Eq. (4) remains the same, and our conclusions about the best-fitting function *I*(*x*) for each prior will remain the same. We find it plausible that some amount of noise perturbs the decision, and not only the estimation of the numbers (see Refs. [6, 31]). But as introducing decision noise would not change our conclusions, we have focused, in the work reported above, on estimation noise, and on how it adapts efficiently to the prior.

The idea that the nervous system encodes stimuli efficiently was first proposed by H. Barlow [1], and finds a modern formulation in the recent literature on neural coding [5, 23–25, 28, 32]. Previous studies have provided experimental evidence that the encoding of stimuli is efficiently adjusted to the stimuli statistics [33, 24, 19, 30, 21, 5, 26]. With the exception of the ‘contextual modulation’ presented in Ref. [26], a common assumption is that the encoding strategy is permanently adapted to some unchanging distribution of stimuli, encountered in nature over long timescales. Here we show, conversely, that subjects are able to adapt their encoding over a relatively short time-scale, the one-hour timeframe of the experiment (an adaptation to a change in the prior, within the timeframe an experimental session, is also found in Ref. [34]). This raises the question, which we leave for future investigations, as to how the brain dynamically tunes its representational capacity to changing environmental statistics.

Our study furthermore differs from these psychophysics results in that the stimuli presented are Arabic numerals. The variables relevant for decision are thus not physical magnitudes (such as the lengths of bars), but magnitudes represented in symbolic form, which one could expect to be processed in a different way (e.g., ‘25’ and ‘52’ have the same size and luminosity, but carry different semantic meaning). Studies on numerosity perception, however, suggest that there are important analogies with sensory perception [35–39]. For instance, subjects exhibit scalar variability in various numerosity tasks [40–42], which has also been interpreted as resulting from an efficient representation of numbers, adapted to a ‘natural’ number distribution [43]. Our results supports the hypothesis that number processing can be understood using the same theoretical framework as accounts for sensory perception. Likewise, Ref. [44] have used this framework to account for the subjective valuation of food items. In addition, we show that the encoding-decoding of numerals can quickly adapt to positively and negatively skewed distributions of numbers.

The efficient-coding, Bayesian-decoding model we examine predicts that estimates should be biased, and that a simple mathematical relation exists between the bias and the derivative of the estimation variance (Eq. (4)). A quantity related to the estimation variance and often measured in perceptual tasks is the discrimination threshold, *DT* (*x*), which quantifies the sensitivity of an observer to small changes in the stimulus. From Eq. (4) we can derive, as shown in Methods, a relation between the bias and the discrimination threshold, as

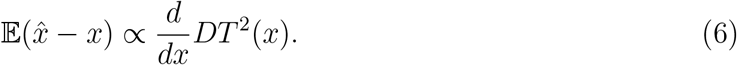

Wei and Stocker [26] derive a relationship of this form, but under an assumption that the encoding maximizes the mutual information between the number and its internal representation. They show that it is supported by numerous empirical results obtained in perceptual tasks, and thus they call it a “law of human perception”. Morais and Pillow [28] obtain this relation when encoding solves a Fisher-information optimization problem like the one that we consider (Eq. (3)), but assuming that estimates are given by a maximum-a-posteriori (MAP) estimator, while we assume a Bayesian-mean estimator. Moreover, in the version of Eq. (6) that they derive from the MAP estimator, the constant of proportionality is necessarily negative in sign, whereas a positive sign is implied by the Bayesian-mean estimator (when *p* < 1), and this is required in order to fit our empirical results.

An assumption of the efficient-coding, Bayesian-decoding model is that subjects hold a subjective prior about the numbers. If the subjects’ prior matched the actual distribution of numbers used in each of the three prior conditions, the noise function, *s*(*x*), should equal the inverse prior raised to the exponent 1*/*(*p* + 1). Although in the Downward and Upward conditions the two functions are qualitatively comparable, discrepancies remain; and in the Uniform condition, the prior is flat while the noise function is U-shaped, and larger for large numbers (Fig. 4). A possible explanation is that subjects may not hold the correct prior. They might, for instance, start the experiment with a prior that corresponds to the frequencies of numbers that one usually encounters, and not fully update these prior beliefs during the task, in spite of our efforts to familiarize them with the correct distribution (see Methods). The frequency of numeric quantities typically observed follows approximately a power law [43, 45, 46], i.e., a distribution skewed towards small numbers. In the Uniform condition, if the subjective prior is in-between the natural, power-law prior, and the correct, Uniform prior, it should also be skewed, to some extent, towards small numbers. The efficient noise function, in this case, would be larger for large numbers, which is precisely what we observe in subjects’ data (Fig. 4). Incorrect subjective priors are thus a plausible source of the observed discrepancies between the subjects’ noise functions and those predicted by the correct priors.

A deviation of the noise functions away from theoretical predictions is found near the endpoints of the interval of presented numbers: the noise increases, despite the rising prior density, near the lower endpoint in the Downward condition, and near the upper endpoint in the Upward condition. The model we investigate relies on a small-noise assumption (in particular, the quantity minimized in Eq. (3) is a lower bound on the mean *L*_*p*_ error only in the limit of a vanishing noise [28]). When noise is large, the optimal Fisher information differs from a simple power of the prior — in particular at the boundaries [25], if the mutual information is maximized (*p* → 0). The optimal Fisher information, for a number *x*, presumably does not depend only on the local value of the prior at this number, *π*(*x*), but depends instead on a larger neighborhood around it. This should distort the small-noise solution, especially at the endpoints where the prior vanishes abruptly after reaching its highest value. Subjects’ data seem consistent with this account, but it calls for further theoretical investigation away from the small-noise hypothesis.

A different account of the observed discrepancies is that the subjects do not apply exactly the efficient-coding optimization program we assume, in which a lower bound on the mean *L*_*p*_ error is minimized (or an approximation of the mutual information is maximized, in the limiting case *p* → 0; Eq. (3)). In our task, subjects might instead maximize the financial reward they can expect; the optimal Fisher information is presumably somewhat different for a different objective. An alternative hypothesis is that subjects adapt their encoding strategy to the prior, but choose to encode with higher precision the numbers that are close to the prior mean, instead of optimizing the program described by Eq. (3). The three bestfitting noise functions, indeed, reach their minima near the priors’ means (Figs. 3a, 5a). This could be a general strategy, used for stimulus encoding in various situations, or it could stem from the specifics of our tasks, in which subjects are asked to compare empirical averages. In any case, we note that the proportionality relation between the bias and the derivative of the noise variance (Eq. (4)) relies on the choice of a noise function that is efficient in the sense of our optimization program (Eq. (3)). A different choice of encoding, such as one that minimizes the noise near the mean, would result in a different relation (if any), and presumably not one of proportionality. However, when fitting the bias and the noise separately, we find that they are consistent with the proportionality relation (Fig. 3b); and the model that directly implies this relation is our best-fitting model. (Besides, in sensory domains, the proportionality relation is supported by many experimental results, as mentioned above.) We cannot, however, reject this hypothesis; it calls for finer investigations on the influence of task reward and prior distribution on the choice of an encoding strategy.

## Methods

### Experiment, subjects, and reward

37 subjects, 18 female and 19 male, aged 24.5 on average (s.d.: 8.6), participated in the experiment. All but one subject participated in two blocks of trials, in each of which all numbers were samples from a single distribution (Uniform, Upward or Downward), so that each subject (except one) experienced two of the three conditions. Each session lasted about one hour. Subjects were explicitly told the current distribution, and at the beginning of each new block they were presented with a series of random samples, in order to familiarize them with the distribution. On screen, the numbers were presented with two decimal digits, and for a duration of 500ms. In each trial, the color of the first presented number was chosen between red and green with equal probability. The score of the subject was augmented at each trial by the chosen average, and thus increased over the course of the experiment. At the end of the experiment, the subject received a financial reward, which is a linear function of the total score, with a $10 “show-up” minimum. The expected reward, not taking into account the $10 minimum, for a hypothetical subject providing random responses (i.e., choosing “red” with probability 0.5 at all trials) was $10, and an accuracy of 80% of correct responses yielded an average of $25. The average reward over the 37 subjects was $28 (s.d.: 3.8). 33 subjects experienced two blocks of 200 trials (so that 2×200=400 decisions were collected per subject), three subjects experienced two blocks of 210 trials, and one subject experienced one block of 210 trials (totaling 14,670 trials). Three trials were excluded from the analysis because the number 100.00 erroneously appeared in these trials. A total of 4610 responses were collected in the Downward condition, 5040 in the Uniform condition, and 5017 in the Upward condition. All participants provided informed consent. The procedures of this experiment comply with the relevant ethical regulations and were approved by the Institutional Review Board of Columbia University (Protocol Number: IRB-AAAR9375). The task was coded in the Python programming language and with the help of the PsychoPy library [47].

### The transformation *m*(*x*) and decision weights

To shed light on the relation between the transformation *m*(*x*) and the decision weights, we derive an approximation to the probability *P* (*red* | *x*) of choosing ‘red’ conditional on a red number *x* being presented. Consider first the model with constant noise (*N*(*m*(*x*), *s*^2^)). The probability can be computed by marginalization of the probability of choosing ‘red’ conditional on ten numbers as

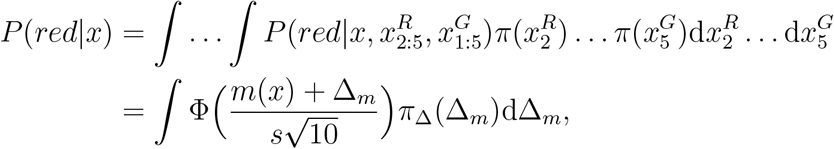

where

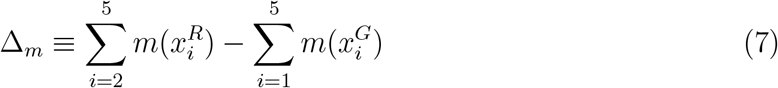

and *π*_Δ_ is the prior density for this random quantity. We approximate *π*_Δ_ by a Gaussian distribution with the same mean and variance, so that

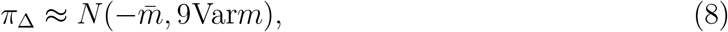

where 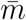 is mean of *m*(*x*) under the prior and Var*m* is the variance. Substituting this approximation in Eq. (7) results in

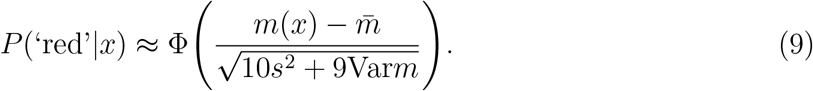

We find this approximation to be fairly close to the conditional probabilities obtained through simulations of the model. In case of variable noise (*N*(*m*(*x*), *s*^2^(*x*))), we replace in Eq. (9) the noise variance *s*^2^ by its average under the prior, 𝔼*s*^2^(*x*), and despite this coarse approximation, we also obtain a close match to simulated data. Figure 5d provides an example of the quality of these approximations, in the case of the efficient-coding, Bayesian-decoding model.

Equation (9) implies that the decision weight, |*P* (‘red’|*x*) −0.5 |, vanishes at approximately the number whose transformation equals the average transformation. In the absence of bias (*m*(*x*) = *x*), this number would be the prior mean; but it is slightly greater in the case of the fitted transformations. Hence the decision weights at the prior mean do not fall to zero, and consequently numbers above and below the prior mean have different weights (Fig. 2c).

### Model fitting and identification

When fitting subjects’ data to the general one-stage model *N* (*m*(*x*), *s*^2^(*x*)), it is important to note that the transformation *m*(*x*) can be identified from choice data only up to an arbitrary affine transformation: a model with transformation *αm*(*x*) + *β* and noise *αs*(*x*) makes the same predictions as the model with transformation *m*(*x*) and noise *s*(*x*), for any non-zero *α* and any *β* (see Eq. (2)). In calculating the bias plotted in Figure 3b, we choose the ‘scale and location’ parameters *α* and *β* to satisfy two additional desiderata. First, for any choice of *α*, we choose *β* so that the left- and right-hand sides of Eq. (4) are exactly equal at the particular value of *x* where the right-hand side is equal to zero. And second, given this, we choose *α* so as to minimize

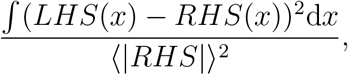

a measure of the relative difference between the two functions *LHS*(*x*) and *RHS*(*x*) defined by the two sides of the equation.

### Behaviour captured only by models with both transformation and varying noise

Here we exhibit patterns in the behaviour of subjects that are not captured by the one-stage noisy-estimation model with general transformation *m*(*x*) but constant noise *s, N* (*m*(*x*), *s*^2^), but that can be captured by a model with both a nonlinear transformation and variable noise, *N* (*m*(*x*), *s*^2^(*x*)), or by the efficient-coding, Bayesian-decoding model. The choice probability implied by the constant-noise model can be written as Φ(*z*), where *z* is

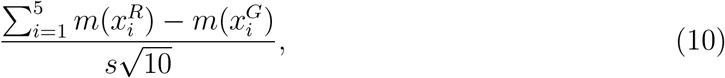

whereas the choice probability implied by a model that additionally features a variable noise 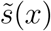 is given by Φ(*z*) with *z* now defined more generally as

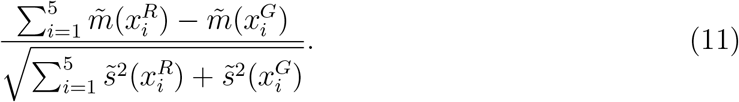

Thus, over a set of presented numbers 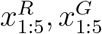 that all imply the same value for the numerator of *z*, the predicted choice probability in the constant-noise case is constant, while it varies with the values of the presented numbers in the variable-noise case. Thus an examination of the choice probabilities of the subjects over different sets of presented numbers satisfying the constraint 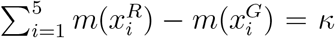, for some constant *κ*, should allow us to discriminate between the predictions of these different model classes.

However, we observe subjects’ behaviour for only a finite number of points 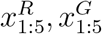 in the space of [10.00, 99.99]^10^ possibilities, and thus for any value of *κ* we have at best a few points verifying this constraint. We therefore choose a coarser version of the constraint:

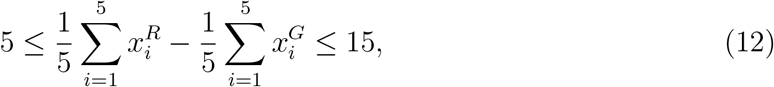

i.e., we consider the trials in which the difference in the averages of the presented numbers is between 5 and 15. We look at how the choice probabilities in these trials varies as a function of the average value of the ten numbers,

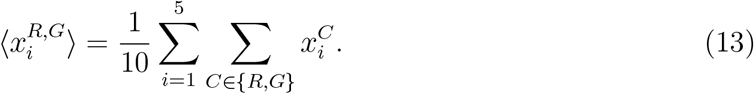

In short, we examine, for trials in which the difference in the averages is around 10, how the choice probability depends on the overall average magnitude of the presented numbers. We find that in the Uniform and Downward conditions, subjects’ choice probabilities decrease as a function of the overall average, whereas they increase in the Upward condition (Fig. 6a). In other words, in the Uniform and Downward conditions the choice probability is closer to 0.5 (incorrect decisions are more frequent) when the presented numbers are large, than when they are small, while the opposite occurs in the Upward condition.

**Figure 6:**
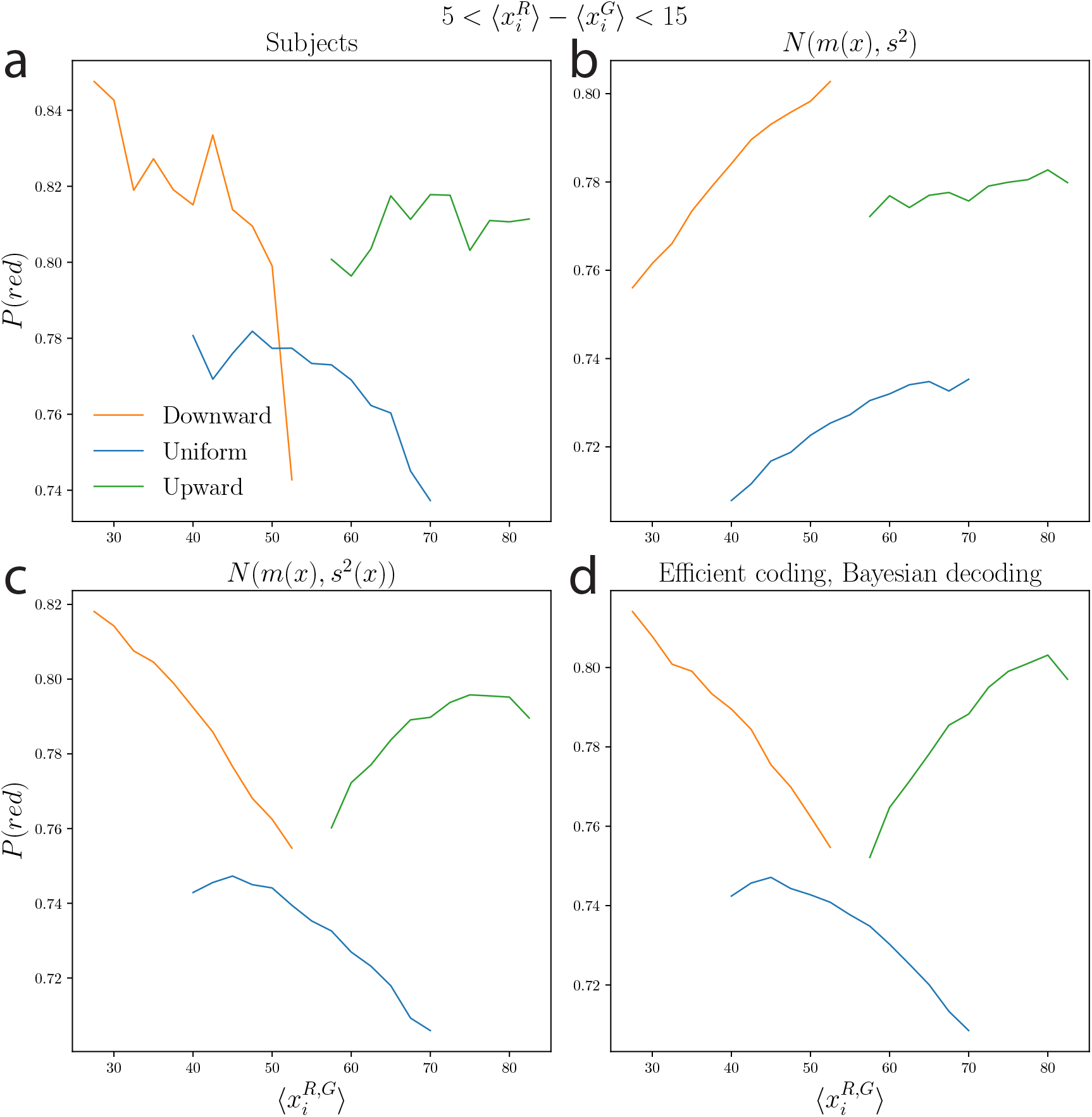
Choice probability as a function of overall average magnitude of numbers. Considering only the trials in which the constraint (12) is verified, the probability of choosing ‘red’, as a function of the average (13) of the ten numbers presented, for subjects (**a**), for the model with a transformation *m*(*x*) but constant noise (*N*(*m*(*x*), *s*^2^), **b**), for the model with a transformation and variable noise (*N*(*m*(*x*), *s*^2^(*x*)), **c**), and for the efficient-coding, Bayesian-decoding model (**d**). All analyses use the subjects’ combined data.

A model with a nonlinear transformation *m*(*x*) but constant noise *s* does not capture this behaviour: in this kind of model, the choice probability is predicted to increase as a function of the overall average in all three conditions (Fig. 6b). In contrast, the model with both a transformation and variable noise (*N*(*m*(*x*), *s*^2^(*x*))), and the efficient-coding, Bayesian-decoding model (which implies a particular kind of variable noise), exhibit the same patterns in behaviour as those of the subjects: the choice probability increases in the Upward condition, and decreases in the Uniform and Downward conditions (Fig. 6c,d). In these latter two conditions, the noise *s*(*x*) at large numbers is appreciably larger than at small numbers (see Fig. 5a, main text), which results in choice probabilities closer to 0.5 at large numbers. These results support the hypothesis that the models which feature both a transformation and variable noise capture more accurately the behaviour of subjects, as also indicated by the model comparison statistics reported in the main text.

### Bayesian model selection using individual data

We conduct a model-fitting and model-selection procedure which differs from that used in the main text in two ways. First, cross-validation, instead of the Bayesian Information Criterion, is used to evaluate the likelihoods of the models while penalizing model complexity. Second, we implement a Bayesian random-effects analysis, which takes subjects heterogeneity into account.

For each subject and each of our 10 models (five estimate models, with either homogeneous or prior-specific parameters), we estimate the likelihood of the model, for that subject, through 10-fold cross-validation. Specifically, we split the subject’s data in 10 interleaved subsets. For each subset, we fit the model by maximizing the likelihood of the 9 complementary subsets, and then use the best-fitting parameters to compute the ‘out-of-sample’ likelihood of the subset. With this procedure, for 83.8% of subjects, the highest total likelihood is obtained with the efficient-coding, Bayesian-decoding model, in which the bias is proportional to the derivative of the variance. 13.5% of subjects are best fitted by the model with transformation of the number, and constant noise (*N*(*m*(*x*), *s*^2^)). Only 2.7% of subjects are best fitted by the model with both transformation and varying noise (*N*(*m*(*x*), *s*^2^(*x*))), in spite of the absence of constraints relating these two functions, in this model. Furthermore, for 75.7% of subjects the highest total likelihood is obtained with prior-specific parameters, i.e., when the parameters of the model are allowed to depend on the prior condition (except the parameter *p*, in the efficient-coding, Bayesian-decoding model).

The results just presented relate to which model yields the largest likelihood, for each subject, but neglect the relative values of the likelihoods of the various models, and thus provide no indication on the weight of the evidence favoring one model over another. We conduct a finer analysis through the ‘Bayesian model selection’ (BMS) procedure described in Ref. [27]. This method assumes that there is a multinomial distribution over the 10 models, parameterized by the vector of probabilities of the models, 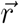, and that the behaviour of each subject is randomly chosen among the 10 models, according to this distribution. The vector 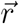 is unknown, but the BMS method prescribes a way to obtain a posterior over the vector 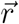, as a Dirichlet distribution, parameterized by a vector 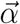, of size equal to that of 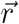.

We implement this procedure and obtain the parameters 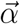 of the Dirichlet posterior distribution. Following the literature, we report, in Table 1, the parameters 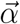, the expected probabilities 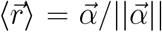of the models, and the ‘exceedance probabilities’, i.e., the probability that each model is more likely than the other models. The expected probability of the efficient-coding, Bayesian-decoding model is substantially larger than the other models: 68.8%, with prior-specific parameters, and 10.6% with homogeneous parameters. One advantage of the BMS method is that it allows to compute the expected probability of hypotheses that correspond to classes of models; for instance, in our case, the expected probability that the parameters are prior-specific, by simply adding the expected probabilities of the corresponding models. We find that the expected probability that the parameters depend on the prior is 80.9%, thus substantially larger than the probability that they are homogeneous across prior conditions. Finally, we compute numerically the exceedance probabilities, through Monte-Carlo estimation, using 10 million samples from the Dirichlet distribution. Out of these samples, in eight cases the most likely model was the efficient-coding, Bayesian-decoding model with homogeneous parameters, and for all the other samples the most likely model was the efficient-coding, Bayesian-decoding model with prior-specific parameters.

**Table 1:** Bayesian Model Selection favors the efficient-coding, Bayesian-decoding model with prior-specific parameters. Parameter vector 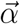 of the Dirichlet posterior distribution over the multinomial distribution of the ten models, computed following Ref. [27], along with the implied expected probabilities of the multinomial distribution and the exceedance probabilities.

| Model | Parameters | Dirichlet $\vec{\alpha}$ | Expected prob. $\langle \vec{r} \rangle$ | Exceedance prob. |
| --- | --- | --- | --- | --- |
| $N(x, s^2)$ | Homogeneous | 1.00 | 2.13% | 0% |
|  | Prior-specific | 1.00 | 2.14% | 0% |
| $N(x, s^2(x))$ | Homogeneous | 1.00 | 2.13% | 0% |
|  | Prior-specific | 1.00 | 2.13% | 0% |
| $N(m(x), s^2)$ | Homogeneous | 1.00 | 2.13% | 0% |
|  | Prior-specific | 2.60 | 5.51% | 0% |
| $N(m(x), s^2(x))$ | Homogeneous | 1.01 | 2.16% | 0% |
|  | Prior-specific | 1.06 | 2.26% | 0% |
| Efficient coding,<br>Bayesian decoding | Homogeneous | 4.97 | 10.58% | .00008% |
|  | Prior-specific | 32.35 | 68.82% | 99.99992% |

Figure 7 shows the best-fitting noise functions for each subject, in the three prior conditions. Consistently with the functions fitted on pooled data, presented in the main text, for a majority of subjects the noise function is broadly increasing in the Downward-prior condition, and decreasing in the Upward-prior condition. More quantitatively, for instance, in the Upward-prior condition, the average value of the noise function *s*(*x*) over the first half-interval (from 10 to 54.995) is greater than its average value over the second half-interval (from 54.995 to 99.99) for 80.0% of subjects, i.e., the encoding of a large majority of subjects is more noisy for the less frequent numbers. In the Downward-prior condition, consistently, the average value of the noise over the first half-interval is lower (higher precision) than the average noise over the second half-interval, for 78.3% of subjects; and in the Uniform condition, for 72.0% of subjects. Finally, the best-fitting values, across subjects, of the parameter *p* (median: 0.74, mean: 0.60, s.d.: 0.49) are consistent with the value obtained with the pooled data (0.70).

**Figure 7:**
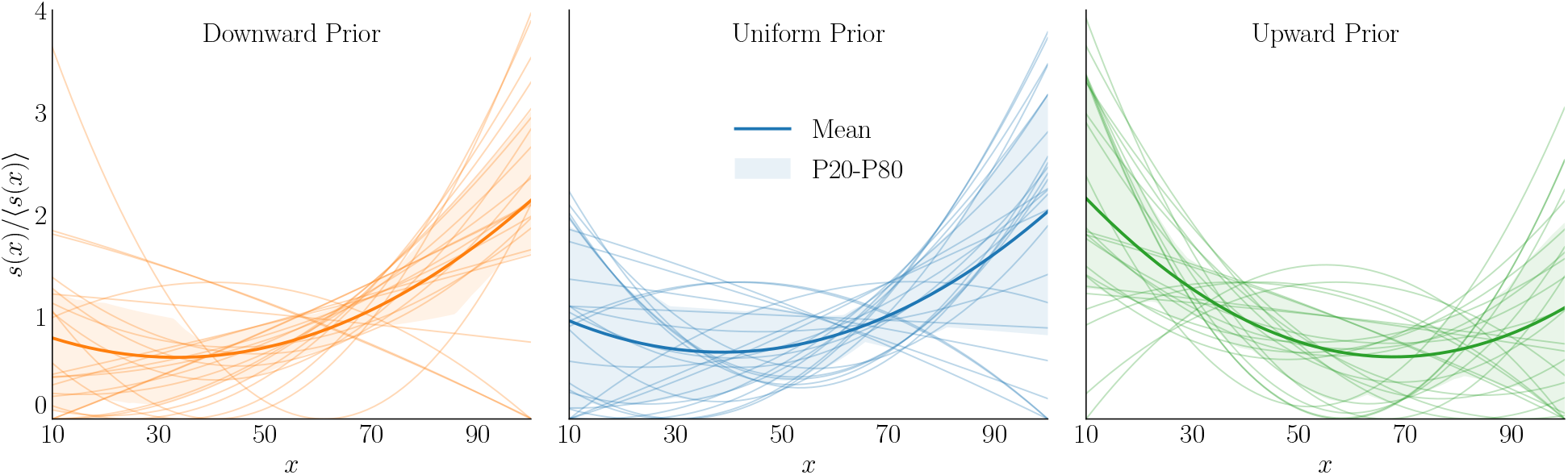
Subjects’ noise functions are adapted to the prior distributions. For each subject and in each prior condition (Downward, *left*, Uniform, *center*, and Upward, *right*), best-fitting noise function of the subject, *s*(*x*), divided by its average over the interval of numbers, ⟨*s*(*x*)⟩, so as to obtain comparable curves across subjects (thin lines); mean function across subjects (thick lines); and 20th and 80th percentiles (shaded areas).

The results of our investigation on the individual behavior of subjects strengthen our main conclusions. First, the responses of a large majority of subjects are better captured by models with prior-specific parameters, rather than parameters that are homogeneous across priors, suggesting that the way subjects encode the presented numbers depends on the statistics of the numbers in each condition — and that the subjects were able to change their encoding within the duration of the experiment (one hour). Second, the subjects’ best-fitting noise functions, in the Upward and Downward prior conditions, differ in the way that our model of efficient coding and Bayesian decoding would predict, if we suppose that the subjective priors for which encoding and decoding are optimized reflect at least partial learning of the differing frequency distributions in these conditions. In each case, subjects encode the (objectively) less likely numbers with more noise, as should happen if the encoding is optimized for subjective priors that assign more probability to high numbers in the Upward condition and more probability to low numbers in the Downward condition. Third, the best-fitting model for most subjects is the efficient-coding, Bayesian-decoding model, in which we require that the subject’s bias and variance functions for each condition be consistent with the theoretical restriction stated in Eq. (4). This means that we require that encoding and decoding both be optimized for a subjective prior, though we do not require the subjective prior to coincide with each condition’s objective prior. The relative success of this more parsimonious model suggests that the “law of human perception” proposed by Wei and Stocker [26] applies not only to sensory perception, but also to the understanding of quantities presented symbolically.

### Optimal Fisher information

The Fisher information function *I*(*x*) that solves the optimum problem defined in Eq. (3) is obtained through variational calculus. Let *J* be the functional

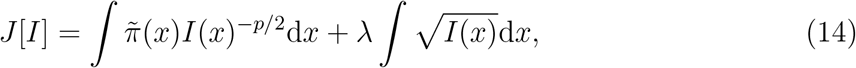

where *λ* is the Lagrange multiplier associated with the encoding constraint in Eq. (3). Taking the functional derivative with respect to *I* and setting to zero, we obtain

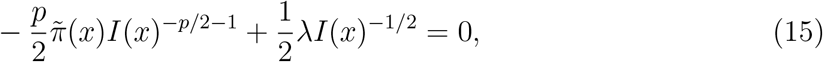

and thus

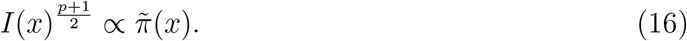

### Efficient-coding, Bayesian-decoding model: relations between bias, variance, and Fisher information, and Cramér-Rao bound

Reference [29] shows that with the assumed form of encoding (a series of *n* independent signals, characterized by a total Fisher information *I*(*x*)), and with a Bayesian-mean decoding that uses the prior 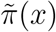, the bias of the decoder is asymptotically approximated by

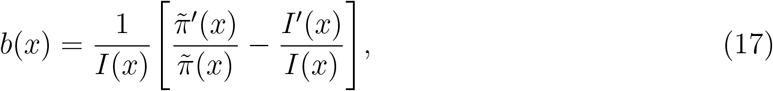

where 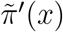 and *I*′(*x*) are the derivatives of 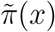 and *I*(*x*). The Fisher information is of order *n*, thus the bias is of order 1*/n*. As for the variance, it is asymptotically approximated by 1*/I*(*x*); this approximation is accurate to order 1*/n*. We use these approximations to define the bias and the variance of the efficient-coding, Bayesian-decoding model. Regarding the variance, the Cramér-Rao lower bound on the variance of biased estimators is

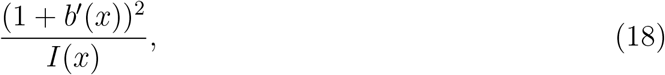

where *b*′(*x*) is the derivative of the bias. Because Equation (17) implies that the bias *b*(*x*) is of order 1*/n*, this bound is equal to 1*/I*(*x*), to order 1*/n*. Note that to order 1*/n* this is the variance of the Bayesian posterior mean decoder, as reported in Eq. (5). Thus the difference between the Cramér-Rao bound and the variance of the estimates, in our model, is within the magnitude of our approximation errors, and asymptotically vanishing.

As for the bias (Eq. (4)), the choice of the Fisher information (see Eq. (16) above) implies that

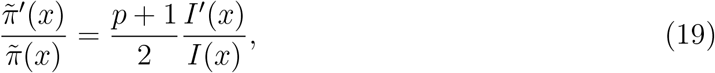

and thus the implied bias is

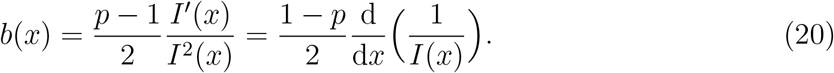

### Large biases, and their overestimations due to the asymptotic approximation

The magnitude of the bias of our best-fitting model, as shown in Fig. 5b, might seem implausibly large, with biases greater than +100 in the Downward and Uniform conditions, and below −100 in the Upward condition. This implies that the subjects’ estimate of the average of the presented numbers could sometimes be below 0, or above 100, although all of the numbers are actually positive and below 100. The large biases, however, are predicted only for extreme values of *x* that do not occur very often under those conditions. In fact, the biases are smaller than 50 (in absolute value) 88% of the time in the Downward condition, 78% of the time in the Upward case, and 68% of the time in the Uniform condition. Consequently, it is relatively rare that the estimate of an average of five numbers be extremely biased. Looking at the distribution of the estimates, by the model, of the averages of five numbers, where each number is sampled from the prior, we find that estimates below zero occur in well below 1 percent of all cases: 0.60% in the Uniform condition, 0.25% in the Downward condition and 0.37% in the Upward condition (Fig. 8).

**Figure 8:**
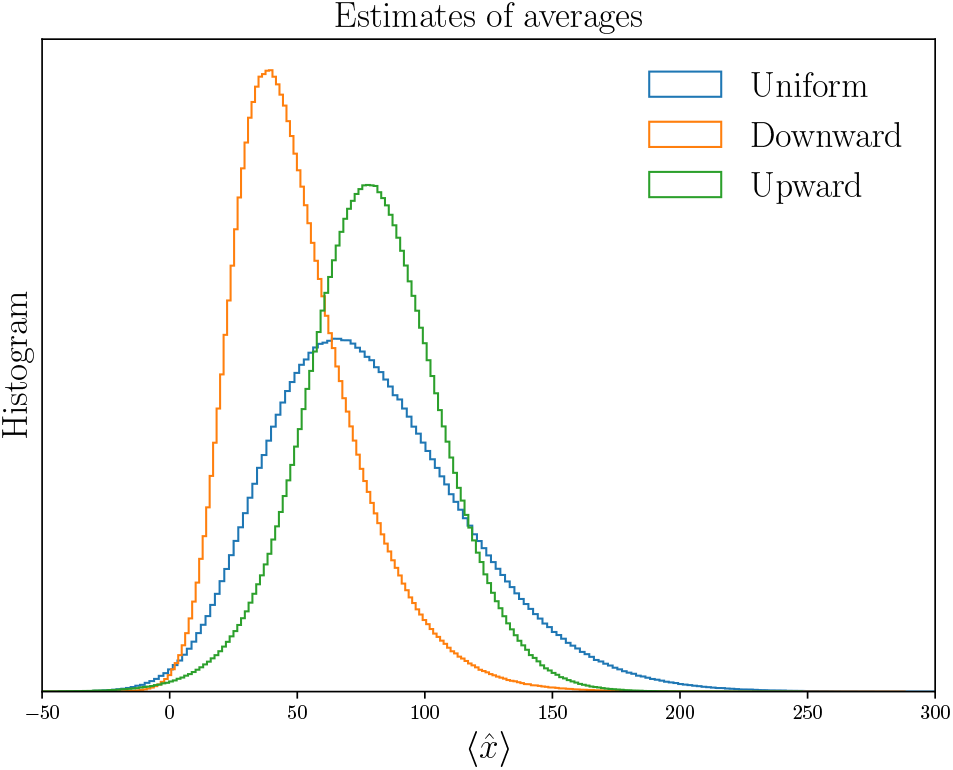
Distribution of the best-fitting model’s estimates of five-number averages. Histograms obtained by sampling from each prior ten million sets of five numbers, and simulating the estimates of their averages by the efficient-coding, Bayesian-decoding model (specified by Eq. (5)).

Furthermore, while a model of Bayesian decoding implies that estimates should always be within the support of the prior distribution, our model does not assume that subjects’ estimates represent Bayesian posterior mean estimates based on the *objectively correct prior* (i.e., the distribution of numbers actually used in the experiment in each condition). Instead, we hypothesize that subjects’ decisions represent optimal Bayesian decisions under an *implicit prior distribution*, that we need to infer from their behaviour. Thus estimates that are occasionally outside the interval [10, 99.99] are perfectly consistent with Bayesian-mean decoding, if we suppose that the subject’s prior has a support wider than the range of numbers actually used in the experiment. Of course, we argue that our subjects’ implicit priors seem to adjust, at least to some extent, to the distribution of numbers actually used; we conclude that the implicit priors are different under the three different conditions, with the mean of the implicit prior highest for the Upward distribution (where the correct prior mean is highest) and lowest for the Downward distribution (where the correct prior mean is lowest). Since we argue that at least some degree of learning appears to occur even over the time scale involved in our experiment, we would be disturbed if the implicit priors required to make sense of the subjects’ behaviour were very different (for example, with a much higher variance) than the actual distributions used in the experiment. But there is nothing implausible about assuming that subjects’ decision rules are optimal for a prior that assigns some probability to the occurrence of numbers more extreme than any of those actually used.

Finally, we note that there is some degree of numerical error involved in the use of our asymptotic (small-noise) approximation in an experimental setting where our estimates imply (and indeed it is reasonable to expect) that the noise in the internal representations of the individual numbers is substantial. Small-noise approximations of the same kind as we use are commonplace in the literature on efficient coding (e.g., Refs. [21, 28]), and the use of this approximation has important advantages: first, of providing analytic expressions for the bias and for the variance of the estimates (Eq. (5)), and second, of allowing to abstract away the specifics of the encoding (such as the detailed properties of encoding neurons). Nonetheless, it is worth considering the errors that this might involve in the present application. It seems likely that our calculations based on the asymptotic approximation exaggerate the size of the bias in the average estimates associated with extreme values of *x*, especially when these are in a region of low prior density (very large numbers, in the Downward condition, or very small numbers, in the Upward condition). In Ref. [29], we show (in the context of a Gaussian prior and a Gaussian encoding) that the error is larger — and the magnitude of the bias is overestimated — for the less frequent stimuli. In our model, the largest biases in the Upward and in the Downward conditions are obtained, respectively, for the small numbers and for the large numbers, i.e., precisely for the less frequent numbers, where the bias is likely to be overestimated.

To illustrate this qualitatively, we run simulations of an encoding-decoding process in which we compute the Bayesian posterior mean estimate taking into account the actual (rather large) degree of noise in the internal representations, rather than relying upon the asymptotic approximation. As in Ref. [29], in these simulations we assume an encoding model with Gaussian noise, i.e., given a stimulus *x* the encoded representation *r* is sampled from a normal distribution with mean *µ*(*x*) and variance *ν*^2^, which are chosen so that the Fisher information of this encoding scheme is precisely that of our best-fitting model (for each of the three conditions). The estimated value 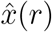 is then computed (numerically) as the mean of the Bayesian posterior. Consistently with our model, the subjective prior 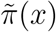 is assumed to be the ‘implicit prior’ for which the Fisher information would be optimal under the asymptotic approximation (i.e., 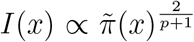). Implementing this proposal requires, however, that we specify the precision of the encoding (i.e., the Fisher information) outside the range of numbers actually used in the experiment. We cannot estimate this from our data, but we do not believe there is any reason to assume that the precision of encoding is zero for all numbers outside the true support (given that the prior need not have been learned too precisely). Thus we choose a Fisher-information function that decreases outside this range as the inverse of the distance to the closest border of the range (i.e., numbers further from the true support are more poorly encoded). These simulations are not meant to constitute a new model (and we do not fit their parameters to behavioral data), but only to give an illustration of the degree to which the bias of a Bayesian decoder in the case of large encoding noise may differ from the bias implied by the asymptotic approximation.

Figure 9 shows the estimates (left panel), and the average biases (right panel) resulting from the simulations of the encoding-decoding process taking into account the degree of noise implied by our estimates (dashed lines) and from the asymptotic approximation used in our model (solid lines). For all three priors, the biases in the two cases follow the same broad pattern: (i) the bias is zero when *x* is equal to the mean of the implicit prior; (ii) the bias is negative for values of *x* below the prior mean, and positive for values of *x* above the prior mean (i.e., the bias is ‘repulsive’); and (iii) the bias increases with *x* at about the same rate on average, over much of the range of values occurring in the experiment. The exact shapes of the bias functions are however different, and the values *m*(*x*) − *x* predicted by the asymptotic approximation are quite different in the case of extreme values of *x* that occur with low frequency (very large values under the Downward condition, very small values under the Upward condition). This suggests that the largest of the biases shown in Fig. 5b are actually artifacts of the inaccuracy of the asymptotic approximation in the case of extreme values of *x*. For small values under the Upward prior, and large values under the Downward prior (i.e., for infrequent values), the asymptotic approximation implies large biases (as shown in Fig. 5b), while the bias in our simulations based on Bayesian decoding for the actual noise variance is more modest; in fact, it is never larger than 30. This suggests that in a more exact calculation of the implications of a Bayesian decoding model fit to our subjects’ data, the biases in subjects’ estimates of individual numbers would also have a maximal magnitude on this order.

**Figure 9:**
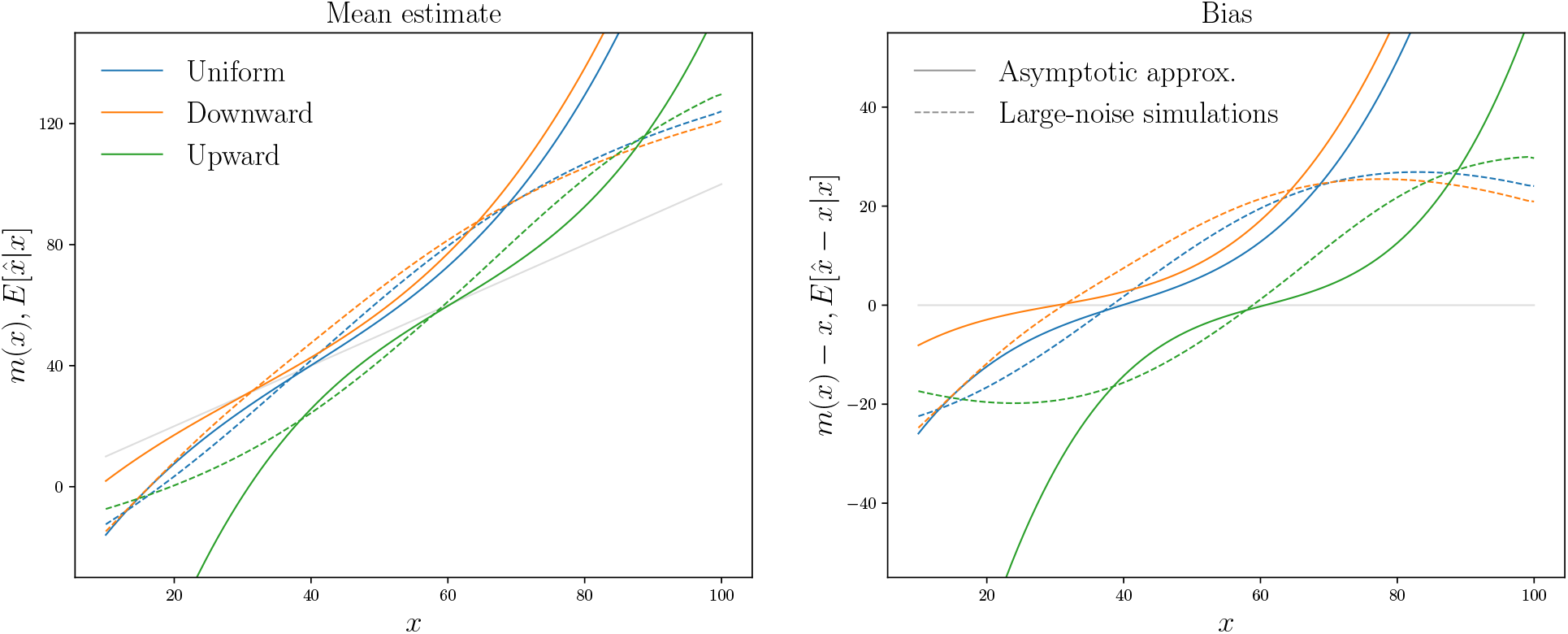
Mean estimate and bias resulting from simulations of an encoding-decoding process, and from the asymptotic approximation. The solid lines in the right panel correspond to Fig. 5b, as our best-fitting model relies on the asymptotic approximation.

One might be concerned that the numerical results in our paper rely upon an approximation that sometimes overestimates the bias in this way; but these overestimations happen infrequently (according to the simulations just discussed), so that overall, they are likely to have only a limited impact on our conclusions (for example, about the Fisher-information function that best fits behavior under each of the three conditions). Specifically, the asymptotic approximation is less accurate when the encoding precision is low; but in our efficient coding model, encoding precision is low precisely for the numbers that are the least frequent (Eq. (16)). As a result, the distributions of implied values for the estimation error, 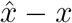, (taking into account the noise in the estimates due to the noise in *r*) are not very different in our simulation using large-noise Bayesian decoding and the calculations using the asymptotic approximation, as shown in Fig. 10. (Note that in this figure the estimation noise is included, while Fig. 9 shows the average bias).

**Figure 10:**
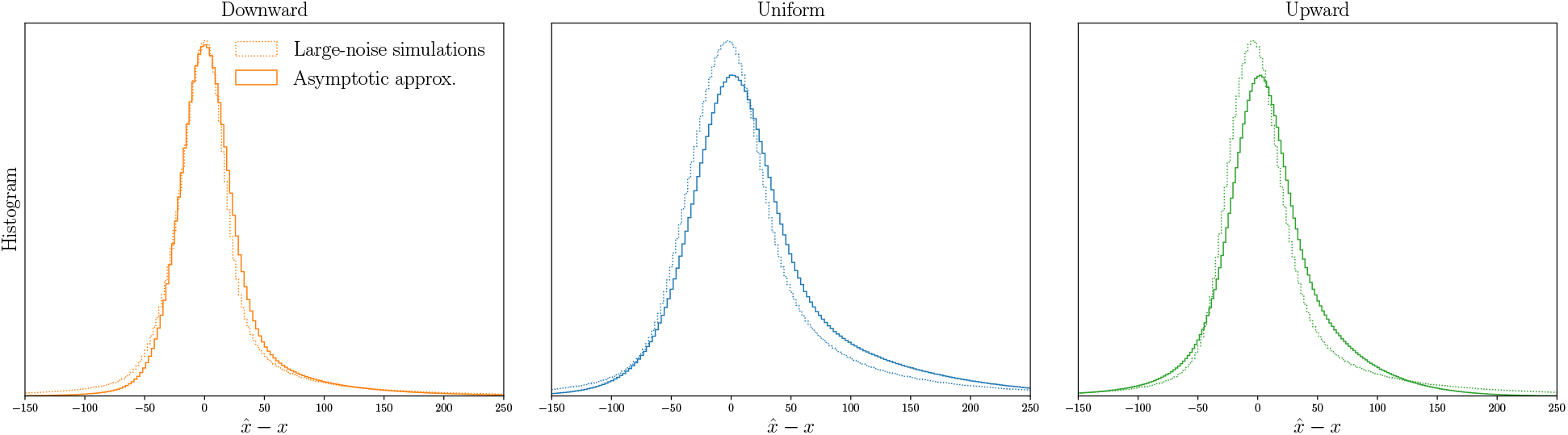
Distributions of estimations errors, 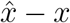, resulting from simulations of an encoding-decoding process, and from the asymptotic approximation.

In summary, the biases in the estimates of numbers by a Bayesian decoder can be somewhat large, because our hypothesis allows for decoding using an ‘implicit prior’ that assigns some probability to the occurrence of values more extreme than those used in the experiment. Moreover, the asymptotic approximation used in our model sometimes overestimates the size of this bias. These overestimations, moreover, are severe mainly in the case of the most infrequent numbers, so that even under the asymptotic approximation, extreme estimates of the average of a sequence of five numbers by our model are very rare, as shown in Fig. 8.

### A different functional form: power function

In the main text, we have chosen polynomials to fit the functions *m*(*x*) and *s*(*x*) with a small number of parameters. Here, we investigate another functional form in which the bias is a sign-conserving power function, similar to the non-linear transformation used in Ref. [6]. More precisely, we study the model implied by Eq. (5), this time assuming that the noise variance, *s*^2^(*x*), is written as

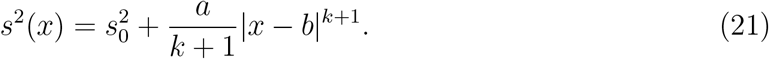

The implied bias (Eq. (4)) is then a sign-conserving power function, namely

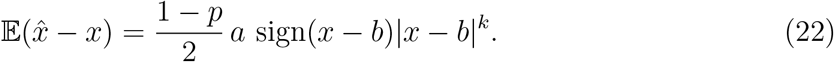

We fit the model with this functional form to our data, and find that its log-likelihood is very close, and slightly higher, than that of the same model fit with a polynomial function (−6750.46 vs. −6751.87). The power-function approach, however, uses four free parameters (*a, b, k*, and 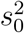), i.e., one more than the polynomial approach (in which *s*(*x*) is a quadratic polynomial). As a result, the BIC with the power function is larger than that with the polynomial function (13612 vs. 13589), so that we conclude that the polynomial form is more parsimonious in its account of the data.

The two functional forms yield very similar results. In the Results section, in order to be consistent with the other models, and for the sake of concision, we present only the results obtained with the polynomial approach. Our results, however, do not depend on a specific choice of functional form.

### Relations between bias, variance, and discrimination threshold

The discrimination threshold *DT* (*x*) is defined as the difference *δ* in stimulus magnitude for which a subject distinguishes two stimuli *x* and *x* + *δ* with a given success rate (e.g., 75%.) In a model in which an estimate 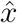 of presented number *x* is normally distributed around a transformation *m*(*x*) of the number, with varying noise *s*(*x*), i.e., 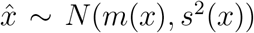, the probability of telling *x*_1_ = *x* and *x*_2_ = *x* + *δ* apart is, assuming *δ* small,

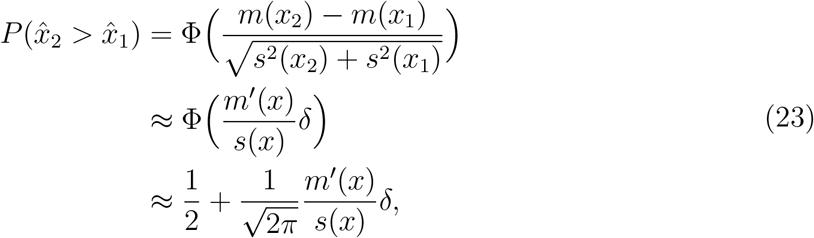

This implies

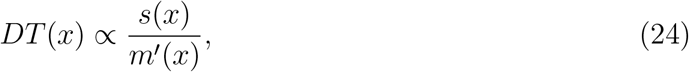

where the proportionality factor depends on the chosen target success rate. In the efficient-coding, Bayesian-decoding model, the bias *m*(*x*) − *x* is proportional to the derivative of the inverse of the Fisher information, therefore the bias is of order 𝒪(1*/n*), and *m*′(*x*) ≈ 1. Equations (24) and (4) then immediately results in Wei and Stocker’s law of human perception [26] (Eq. (6)):

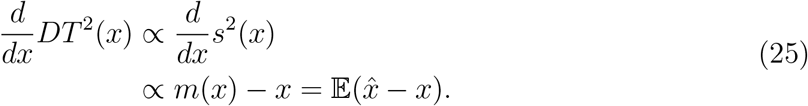

We note, in addition, that the converse derivation appears in the Supporting Information of Ref. [26]: from the relation involving the discrimination threshold (Eq. (6)), the authors derive a relation involving the noise variance, as in Eq. (4).

### A model with variable noise is distinguishable from a model with constant noise

In addition to the results presented above, here we offer a theoretical discussion of the distinguishability of models with constant as opposed to variable noise. We show that, given a sufficient amount of data, a noisy-estimation model with a transformation 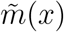 and a variable noise 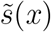 cannot result in the same choice probabilities as a model with transformation *m*(*x*) and *constant* noise *s*, unless 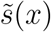 is constant. In this discussion, we assume that 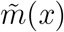, *m*(*x*) and *s*(*x*) are smooth functions, and that 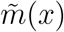 and *m*(*x*) are monotonically increasing. Equality of the choice probabilities implied by the two models requires that

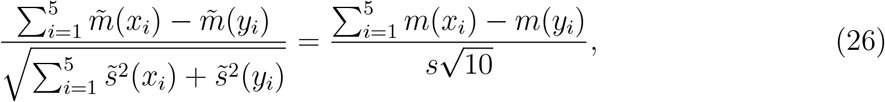

where we denote the red numbers by *x*_*i*_ and the green numbers by *y*_*i*_. Introducing the notation

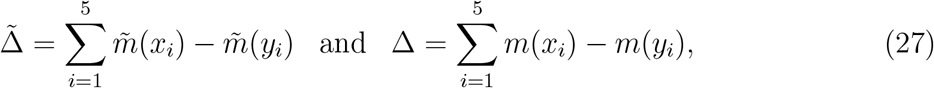

we can rewrite Eq. (26) as

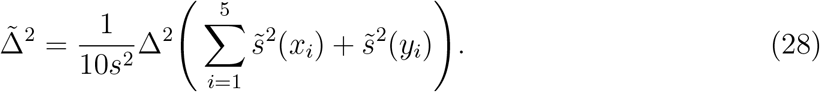

Taking the derivative of Eq. (28) with respect to *x*_*j*_, we obtain

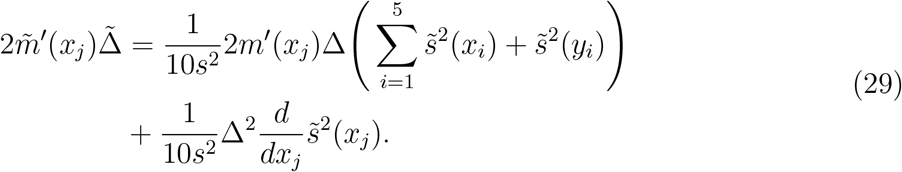

Similarly, taking the derivative with respect to *y*_*k*_, we obtain

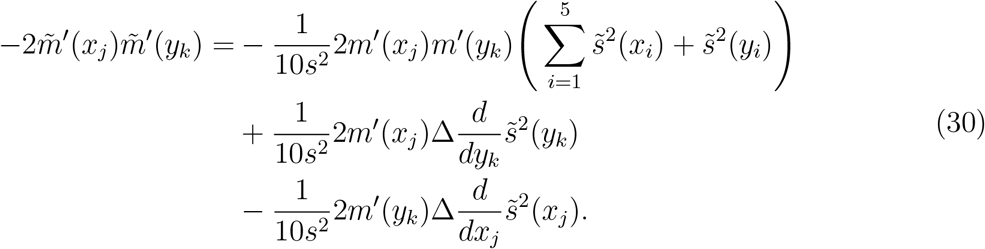

Finally, taking the (third) derivative with respect to *x*_*l*_, we obtain

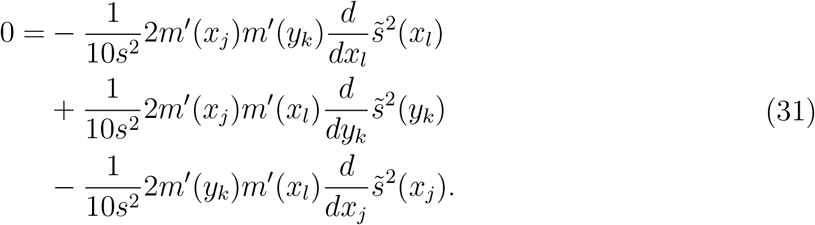

This must be true in particular for *y*_*k*_ = *x*_*l*_. Thus we must have

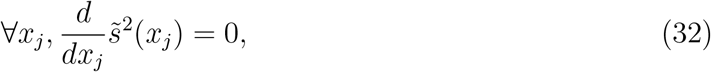

i.e., 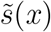 must be a constant function.

## Data availability

Requests for the data can be sent via email to the corresponding author.

## Code availability

Requests for the code used for all analyses can be sent via email to the corresponding author.

## Acknowledgments

We thank Brian Ho for his outstanding help as a research assistant, and the National Science Foundation for research support.

## Author information

### Contributions

M.W. conceptualized the study. M.W. and A.P.C. designed the experiment. A.P.C. implemented the task and collected the data. M.W. and A.P.C. analyzed the data and wrote the computational models. A.P.C. implemented the models. M.W. and A.P.C. interpreted the results and wrote the manuscript.

## Ethics declarations

### Competing interests

The authors declare no competing interests.

## Notes

### Competing Interest Statement

The authors have declared no competing interest.

